# Heparan sulfate modified proteins affect cellular processes central to neurodegeneration and modulate *presenilin* function

**DOI:** 10.1101/2024.01.23.576895

**Authors:** Nicholas Schultheis, Alyssa Connell, Alexander Kapral, Robert J. Becker, Richard Mueller, Shalini Shah, Mackenzie O’Donnell, Matthew Roseman, Weihua Wang, Fei Yin, Ryan Weiss, Scott B. Selleck

## Abstract

Mutations in *presenilin-1 (PSEN1)* are the most common cause of familial, early-onset Alzheimer’s disease (AD), typically producing cognitive deficits in the fourth decade. A variant of *APOE, APOE3 Christchurch (APOE3ch)*, was found associated with protection from both cognitive decline and Tau accumulation in a 70-year-old bearing the disease-causing *PSEN1-E280A* mutation. The amino acid change in ApoE3ch is within the heparan sulfate (HS) binding domain of APOE, and purified APOEch showed dramatically reduced affinity for heparin, a highly sulfated form of HS. The physiological significance of *ApoE3ch* is supported by studies of a mouse bearing a knock-in of this human variant and its effects on microglia reactivity and Aβ-induced Tau deposition. The studies reported here examine the function of heparan sulfate-modified proteoglycans (HSPGs) in cellular and molecular pathways affecting AD-related cell pathology in human cell lines and mouse astrocytes. The mechanisms of HSPG influences on *presenilin-*dependent cell loss and pathology were evaluated in *Drosophila* using knockdown of the presenilin homolog, *Psn*, together with partial loss of function of *sulfateless (sfl)*, a homolog of *NDST1*, a gene specifically affecting HS sulfation. HSPG modulation of autophagy, mitochondrial function, and lipid metabolism were shown to be conserved in cultured human cell lines, *Drosophila*, and mouse astrocytes. RNAi of *Ndst1* reduced intracellular lipid levels in wild-type mouse astrocytes or those expressing humanized variants of *APOE, APOE3*, and *APOE4*. RNA-sequence analysis of human cells deficient in HS synthesis demonstrated effects on the transcriptome governing lipid metabolism, autophagy, and mitochondrial biogenesis and showed significant enrichment in AD susceptibility genes identified by GWAS. Neuron-directed knockdown of *Psn* in *Drosophila* produced cell loss in the brain and behavioral phenotypes, both suppressed by simultaneous reductions in *sfl* mRNA levels. Abnormalities in mitochondria, liposome morphology, and autophagosome-derived structures in animals with *Psn* knockdown were also rescued by simultaneous reduction of *sfl. sfl* knockdown reversed *Psn-*dependent transcript changes in genes affecting lipid transport, metabolism, and monocarboxylate carriers. These findings support the direct involvement of HSPGs in AD pathogenesis.

## Introduction

More than 100 years ago, Alois Alzheimer described a dementia condition associated with a triad of histopathological abnormalities: extracellular deposits of amyloid, intracellular aggregations or neurofibrillary tangles, and accumulation of intracellular lipid in glia, described as “adipose saccules”^1^. These pathological features have guided the strategies to develop a pharmaceutical for Alzheimer’s disease (AD) and amyloid peptide generation and deposition (neuritic plaques) has been one of the principal targets of these efforts. Recently, two antibody pharmaceuticals that reduce the levels of amyloid peptides have been FDA-approved, and one shows demonstrable slowing of cognitive loss^2,3^. Despite this success, there is good reason to identify cellular and molecular pathologies that occur early in the AD process. Further investment in broad, mechanistic approaches to AD development is required to effectively prevent and treat this disease at a level that significantly alters the disease trajectory.

Considerable evidence from a variety of experimental approaches has shown that there are cellular deficits common to AD, implicated in both early, familial AD where causative genetic changes have been identified, as well as late-onset AD, where GWAS studies have found many risk-associated variants that influence AD susceptibility^4–6^. Pathway analyses show enrichment for genes involved in cholesterol metabolism, amyloid plaque and neurofibrillary tangle formation, membrane trafficking, and the innate immune system^4^. Many of the cellular and molecular processes embedded within these gene sets are mechanistically connected^7^. Autophagy, a vital membrane trafficking process compromised in AD, not only removes protein aggregates and damaged organelles (such as mitochondria via mitophagy) but is required for the catabolism of lipids via lipophagy. Mitochondria are also a critical nexus in these mechanisms, providing the catabolic machinery to break down fatty acids via β-oxidation and providing ATP to support cell energetics in the process.

The cellular and molecular events that accompany AD are represented in experimental models of the disease. Many of these are based on human genetic data where specific gene deficits produce early onset, severe AD, and allow detailed examination of the cellular and molecular changes that initiate cell demise^8^. *PSEN1* mutations are the most common cause of familial AD, show dominant inheritance, and produce a cognitive decline in the fourth decade with greater than 90% penetrance^9^. The vast majority (90%) of clinically identified *PSEN1* mutations are loss or reduction in function alleles, reducing the activity of the γ-secretase, suggesting that the dominant inheritance results from reductions in the activity of this critical protein and its associated complex^10^. Recently, the intersection of early and late-onset AD genetics was further highlighted by the discovery of a rare *APOE* variant, *ApoE3 Christchurch (APOE3ch)*, that was associated with the protection of cognitive decline in a 70-year-old with a disease-causing *PSEN1* mutation^11^. Homozygosity for *APOE3ch* in this individual was also associated with reduced accumulation of Tau, while amyloid deposition was unabated and significantly higher than in younger cohorts with cognitive symptoms but lacking the *APOE3ch* variant. The amino acid change in *ApoE3ch* provided a clue as to the mechanism of this variant providing protection against AD symptoms. APOEch bears a change in a lysine residue within the heparan sulfate binding domain of the protein. Biochemical studies with purified APOE proteins demonstrated that APOEch showed dramatically reduced affinity for heparin, a highly sulfated form of heparan sulfate. Furthermore, APOE4, APOE3, and APOE2 proteins showed heparin affinities correlated with their relative risk for developing late-onset AD, with higher affinities associated with greater risk. Finally, a recent report showed that mice expressing a humanized *ApoE3ch* variant showed altered microglial responses and suppression of Aβ-induced Tau seeding and spread^12^. These results suggest that heparan sulfate-modified proteins participate in APOE functions that can modulate AD pathogenesis and influence *PSEN1-*mediated events.

Heparan sulfate-modified proteins or proteoglycans (HSPGs), are abundant molecules of the cell surface and extracellular matrix. The glycan modifications have an important role in their function, serving as binding sites for a wide array of proteins, including growth factors and growth factor receptors. HSPGs modulate the activity of many signaling events, generally serving to increase the activity of secreted protein growth factors. HSPGs are an important class of co-receptors at the cell surface, promoting the assembly of signaling complexes. HSPGs are known to affect signaling via PI3kinase, Akt, and MAPK systems^13,14^. HSPGs regulate Wnt, Hh, FGF, and BMP signaling during development and play a role in both endocytosis and exocytosis of a variety of molecules^15–18^.

We have previously shown that reducing heparan sulfate chain length or sulfation increases autophagy flux in muscle and fat body cells in *Drosophila*^19^. Modest reductions in two critical HS biosynthetic enzyme encoding genes, *sulfateless (sfl)* and *tout velu (ttv)*, either by mutation or RNAi, also rescue cell degeneration in a model of AD mediated by expression of a dominant-negative form of Presenilin in the retina^20^. Cell death and mitochondrial dysmorphology in *parkin* mutants, a model for Parkinson’s, are also suppressed by reducing HS function and rescue depends on an intact autophagy system^20^. Autophagy flux and mitochondrial function, both governed by HSPG activity, are compromised by *PSEN1* deficits^21–24^. In this study, we examine in detail the effects of HSPGs on autophagy, mitochondrial function, and intracellular lipid droplets or liposomes in human cells and mouse astrocytes. Transcriptome changes in human cells defective for HS synthesis reflect the cellular changes in autophagy, mitochondrial biogenesis and lipid synthesis or transport. AD-associated genes identified by large GWAS studies are significantly over-represented in differentially expressed genes in cells lacking HS synthesis. We also show that reducing HSPG-mediated signaling affects mitochondrial function, autophagy, and lipid metabolism in *Drosophila* in a manner that counters deficits produced by *presenilin* knockdown and can suppress cellular phenotypes and molecular signatures in animals with compromised *presenilin*.

## Results

### Transcriptomics of human Hep3B cells defective in HS synthesis shows broad effects on the expression of genes affecting lipid metabolism, autophagy, mitochondria, and AD-associated genes

We have previously reported that reducing heparan sulfate synthesis or modification alters autophagy flux and both function and morphology of mitochondria^19,20,25^. Reducing HS biosynthesis can suppress mitochondrial dysmorphology and muscle cell loss in a fly model of Parkinson’s disease achieved by mutations affecting *parkin*, the fly homolog of human *PARK2*. To determine if HS synthesis serves a conserved role in cellular and molecular events associated with AD pathology, we conducted RNA seq profiling of a human cell line (Hep3B) bearing loss-of-function mutations in *EXT1*^26^, a gene required for HS polymerization. A large number of differentially expressed genes were evident, with 1949 genes showing increased expression and 1294 with decreased expression in *EXT1 -/-* cells (p<0.05, FC≥2.0). We examined genes involved in key cellular processes affected in AD, including autophagy, lipid metabolism, and mitochondria biogenesis and function.

Key components of lipid metabolism are represented in the statin drug pathway (https://www.wikipathways.org/instance/WP430) and provide a broad gene set for assessing differential gene expression affecting lipid. Four gene products in this pathway provide critical functions in the biosynthesis of cholesterol, 1) HMG-CoA reductase (3-hydroxy-3-methyl-glutaryl-coenzyme A reductase (HMGCR), the rate-limiting enzyme in the mevalonate pathway, 2) farnesyl-diphosphate farnesyltransferase 1 (FDFT1), the first specific enzyme in cholesterol biosynthesis, catalyzing the dimerization of two molecules of farnesyl diphosphate in a two-step reaction to form squalene, 3) squalene monooxygenase (also called squalene epoxidase, SQLE), catalyzing the first oxygenation step in sterol biosynthesis and considered one of the rate-limiting enzymes in this pathway and 4) sterol O-acyltransferase 1, a resident enzyme of the endoplasmic reticulum that catalyzes the formation of fatty acid-cholesterol esters (SOAT1). Transcripts encoding all four of these key enzymes are dramatically and significantly reduced in *EXT1* KO Hep3B cells (Supplemental Table 1), suggesting a significant suppression of cholesterol and cholesterol-ester biosynthetic capability. *SREBF2*, encoding a transcription factor that upregulates several genes affecting cholesterol biosynthesis, is significantly downregulated in *EXT1* KO cells. Other genes in this pathway involved in lipid transport/lipoprotein assembly are also significantly downregulated in *EXT1-/-* cells including *APOE, APOC2, APOC1, and APOB*. These proteins are involved in the export of lipids from lipid stores into the bloodstream and tissues. Another gene affecting lipid transport ^19,20^ (MTTP), a protein required for the assembly of β-lipoprotein synthesis and lipid export from cells, is also markedly downregulated in *EXT1-/-* cells.

*APOA4*, encoding a protein promoting the transport of cholesterol from peripheral tissues to HDL, is upregulated in *EXT1-/-* cells. Other lipid transporters with an increased expression upon knockout of *EXT1* include *ABCA7, ABCA10, and ABCA12*. *ABCA7* is notable for the strong linkage of variants in GWAS analysis with AD susceptibility^27,28^, and loss of function alleles are associated with an increased risk of AD in both European and African-descent populations^29,30^. Another gene of interest in lipid biology is *PCSK9*, which encodes a regulator of LDL receptor levels on the cell surface by altering intracellular trafficking. PCSK9 inhibitors are now FDA-approved drugs for lowering LDL levels in plasma. This gene is significantly downregulated in *EXT1* KO cells. MFSD2 lysolipid transporter A (MFSD2A), another gene affecting lipid transport, is dramatically reduced in expression levels in *EXT1* -/- cells. Mice with targeted knockout of *MFSD2A* are leaner and have reduced serum, hepatic, and adipocyte triglyceride levels^31^. It is intriguing that in the brains of the APP/PS1 mice, expression of the proprotein convertase subtilisin/kexin type 9 (PCSK9), and apolipoprotein E (APOE) were all elevated^32^, and both PCSK9 and APOE are reduced in the Hep3B *EXT1* KO cells.

Earlier work showed that reducing heparan sulfate chain elongation or sulfation results in increased autophagy flux to the lysosome in *Drosophila*^19,20^. Genes critical for autophagy function are significantly elevated in *EXT1* KO cells compared to controls, including *SQSTM1, ULK1, PINK1, ATG9B, SRC*, and *PRKAG2*. GSEA of Hep3B *EXT1* KO compared to Hep3B +/+ cells also suggests activation of autophagy. Reactome pathways Selective Autophagy, and Aggrephagy are enriched (FDR=0.24, and 0.25 respectively) and several other related gene sets show similar levels of increased component expression (Autophagy and Lysosome Vesicle Biogenesis) (Supplemental Table 3).

*PPARGC1A*, a master regulator of mitochondrial biogenesis, and *CHD9*, a coactivator of *PPARGC1A*, were significantly increased in *EXT1* -/- cells compared to controls. This is consistent with our data showing increased MitoTracker^TM^ signal in these cells (see below). *PPARGC1A* is required for maintaining normal levels of mitochondrial gene expression and oxidative metabolism^33^. *PPARGC1A* also modulates gluconeogenesis and the capacity for cells to engage in glucose or fatty acid metabolism depending on nutrient availability^34^. *CHD9* is responsible for remodeling chromatin and is a coactivator of *PPARGC1A*^35^.

### Effects of EXT1 knockout on Alzheimer’s Disease Susceptibility Genes

AD is associated with changes in lipid metabolism and transport, and Hep3B cells are derived from hepatocytes, cells critical in these processes. We therefore were interested in differential gene expression profiles in Hep3B *EXT1 -/-* cells affected loci identified as AD susceptibility genes and their relation to functional gene groups implicated in AD pathogenesis. We selected genes identified by GWAS analysis classified at Tier 1 or Tier 2, reflecting different levels of association across 6 GWAS studies performed in populations of European ancestry and published since 2019^36^ (Supplemental Table 2). Overall, 46% of Tier 1 genes showed significant *EXT1-*dependent changes in expression, and Tier 2 representation among the differentially expressed genes was 57%. This reflects a statistically significant enrichment of AD-susceptibility genes among the differentially expressed genes affected by *EXT1 -/-* in Hep3B cells (Chi-square with Yates correction, p<0.0001). g:profiler assessment of the Tier 1 genes affected in *EXT1 -/-* cells showed significant representation of amyloid precursor protein catabolic process (p-adjusted= 6.8x10^-7^). These findings provide evidence for the capacity of heparan sulfate function to preferentially affect genes involved in AD susceptibility.

Large and significant downregulation was found for *FERMT2* and *APOE* (Supplemental Table 2). *APOE* variants represent a substantial genetic risk for LOAD^37^ and the discovery of the *ApoE3ch* variant that both rescues *PSEN1-*mediated EOAD and shows greatly reduced heparin binding suggests an important relationship between heparan sulfate function and AD pathology^11^. *FERMT2* encodes a scaffolding protein affecting integrin and TGF-β signaling and reducing expression in neural stem cells suppresses accumulation of phosphorylated Tau and extracellular APP-derived peptides^38^. Two other genes with differential expression in *EXT1 -/-* cells include *BIN1* and *RIN3. BIN1* and *RIN3* interact directly and together affect Rab5-mediated endocytosis^39^. GWAS analyses have identified both as AD risk loci, and *RIN3* levels have been reported to increase in the hippocampus and cortex of APP/PS1 mice^40^. Elevated expression of *RIN3* in basal forebrain cholinergic neurons (BFCNs) from E16 embryos was associated with enlargement of early endosomes, suggesting an endocytosis abnormality. We have previously reported that compromising heparan sulfate biosynthesis increases activity-dependent endocytosis at the *Drosophila* neuromuscular junction^41^, and recent studies of tissue culture cells with compromised HS synthesis showed increases in endocytosis of GFP-tagged dextran, a measure of clathrin-dependent endocytosis (data not shown). These findings indicated that lowering heparan sulfate-dependent signaling can increase endocytosis, a process that is compromised in AD pathology.

### Conservation of heparan sulfate functions in mitochondrial and lipid metabolism in human cells and mouse astrocytes

We have examined the role of HSPGs in mitochondrial function and lipid metabolism in three different human-transformed cell lines as well as mouse primary astrocytes, derived from either wild-type mice or those bearing humanized variants of APOE that affect AD risk susceptibility in humans. Two independent and previously characterized human cell lines were examined, human melanoma A375 cells bearing CRISPR-mediated knockout of *NDST1*^42^, and a hepatocellular carcinoma cell line, Hep3B, with a directed knockout of *EXT1*^26^ , the cell line used for the transcriptomics analysis described above. Experiments were also carried out with HEK293T cells with lentiviral-mediated knockdown of either *EXT1* or *NDST1.* Mouse astrocytes were isolated from wild-type animals or those bearing humanized *APOE3* or *APOE4* variants^43^. Knockdown of *Ndst1* in these mouse astrocytes was achieved with transient lentiviral expression in culture.

Knockout of *NDST1* in A375 cells has a dramatic effect on the binding and internalization of GNeo, a modified neomycin analog with high affinity and specificity for heparan sulfate^42^. Fluorescent activated cell sorting (FACS) measures of internalized GNeo of A375 *NDST1 -/-* cells compared to wild type A375 showed an approximately 3-fold reduction in internalized GNeo (Figure 1A). Autophagy induction has a profound effect on mitochondria levels and the response of A375 cells to rapamycin, a TOR inhibitor, showed a significant increase in FACS measures of the mitochondrial marker, MitoTracker^TM^ (Figure 1B). These data demonstrate this cell line displays a vigorous autophagy response. The effect of *NDST1*, and hence heparan sulfate structure, on mitochondria was assessed similarly using FACS analysis of MitoTracker^TM^ levels, which increased 1.5-fold, comparable to the effects of rapamycin (Figure 1C). We have previously shown using transmission electron microscopy that A375 *NDST1 -/-* cells have altered mitochondrial morphology^25^, and the MitoTracker^TM^ results support the conclusion that compromising the structure of heparan sulfate has a profound effect on mitochondria. Induction of autophagy is also associated with reductions of intracellular neutral lipids, and we employed FACS analysis of LipidTOX^TM^ staining to determine if the state of heparan sulfate affected lipid metabolism in A375 cells. LipidTOX^TM^ level was significantly reduced in *NDST1 -/-* cells compared to wild-type controls (Fig. 1D).

**Figure 1.**
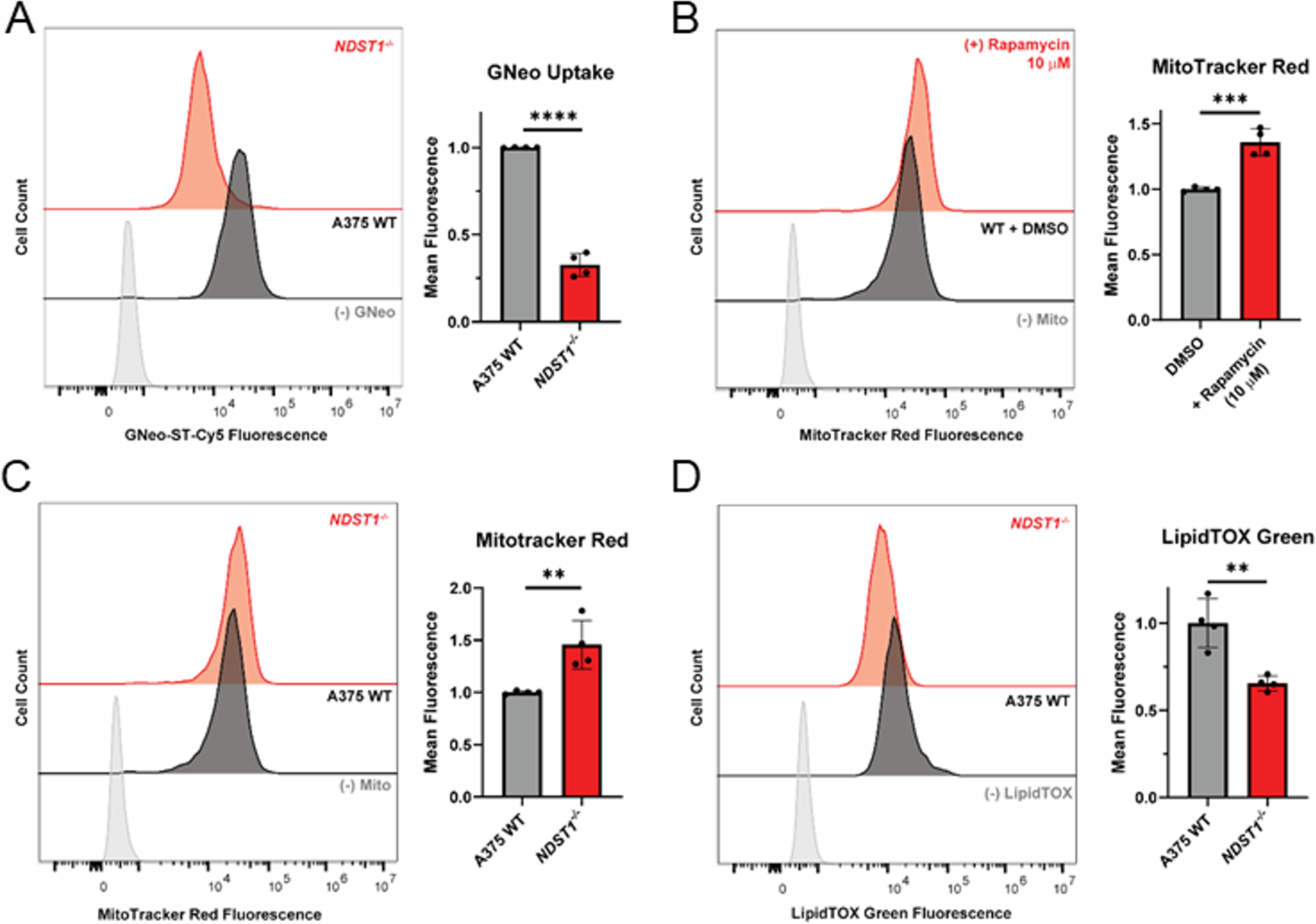
Flow cytometry analysis of A375 human melanoma cells with *NDST1* knockout. (A) Flow cytometry analysis of A375 wildtype and NDST1 knockout mutant (*NDST1^-/-^*) cell lines after 1-hour incubation at 37°C with biotinylated guanidinoneomycin (GNeo-biotin) conjugated to streptavidin-Cy5. NDST1 *N*-deacetylates and adds *N*-sulfate to glucosamine residues of heparan sulfate, which is required for GNeo binding^79^ (t-test, n=4, \*\*\*\**p* < 0.0001). (B) Flow cytometry analysis of A375 wildtype cells after incubation with mTOR inhibitor, rapamycin (10 µM), for 24 hours and subsequent 1-hour incubation with MitoTracker^TM^ Red. Rapamycin significantly increases the MitoTracker^TM^ Red signal (t-test, n=4, \*\*\**p* < 0.001). (C) Flow cytometry analysis of A375 wildtype and *NDST1* mutant cells after 1-hour incubation with MitoTracker^TM^ Red (t-test, n=4, \*\**p* < 0.01). (D) Flow cytometry analysis of A375 wildtype and *NDST1* mutant cells after a 30-minute incubation at 37°C with LipidTox^TM^ Green (t-test, n=4, \*\**p* < 0.01). All error bars represent SEM.

The effect of heparan sulfate biosynthesis on mitochondria was also assessed in Hep3B cells, comparing wild-type to *EXT1* knockout cells. As observed for A375 cells with compromised *NDST1* function, eliminating the activity of EXT1, a chain-elongating heparan sulfate co-polymerase, significantly increased the mitochondrial signal (MitoBlue, Fig. 2A). Metric analysis of confocal images showed significant increases in mitochondrial number and size in *EXT1 -/-* Hep3B cells compared to wild type controls (data not shown).

**Figure 2.**
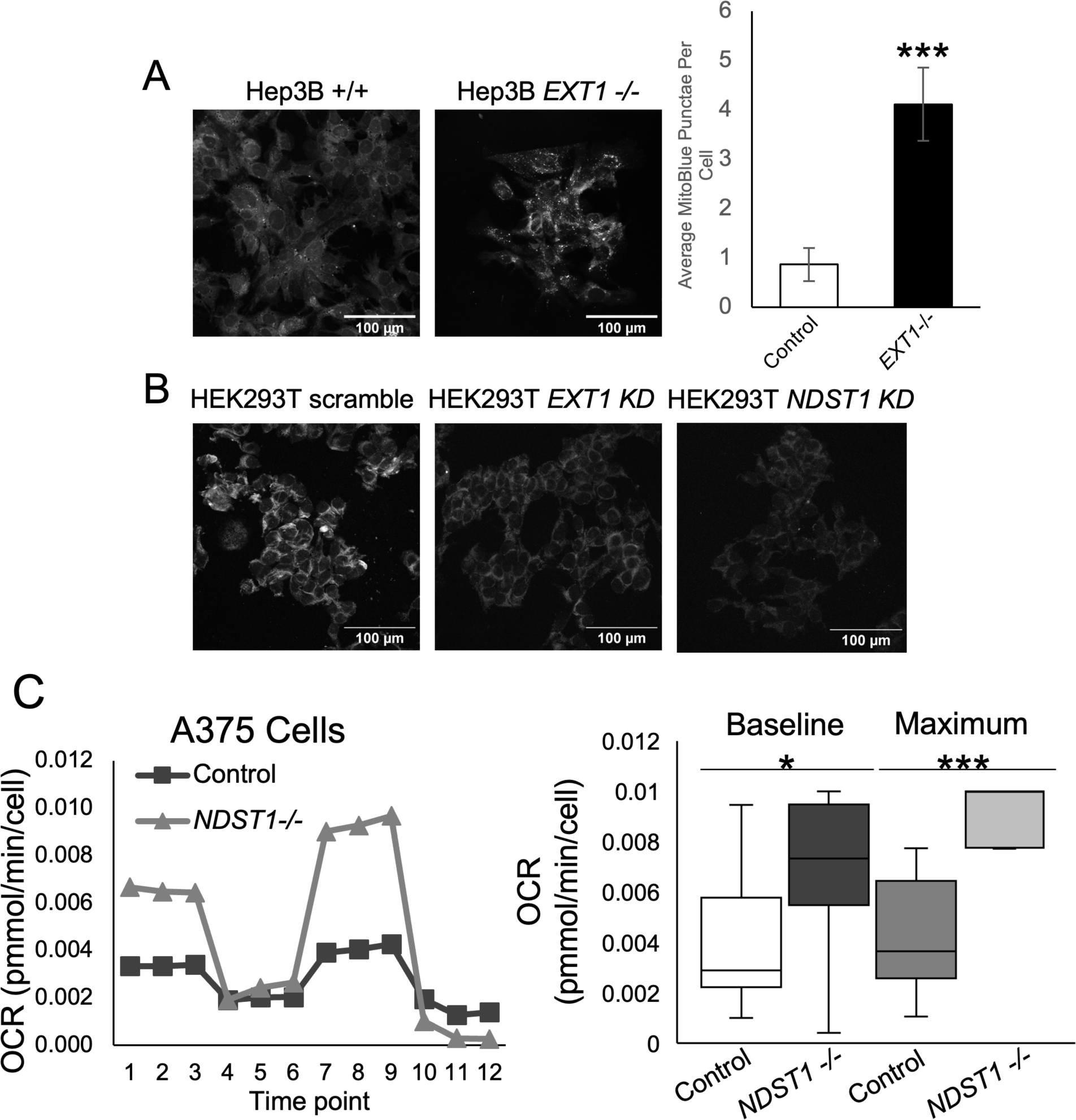
Reduced expression of heparan sulfate biosynthetic enzymes in human cell lines significantly increases the MitoBlue signal, reduces LipidTox^TM^ Red signal, and increases oxygen consumption rate measured by Seahorse Analyzer. (A) Representative confocal fields of Hep3B cells stained with MitoBlue fluorophore. Knockout of *EXT1* significantly increases the number of MitoBlue-positive punctae in Hep3B cells (t-test, Hep3B EXT1 +/+ , n = 17 fields, Hep3B EXT1 -/-, 19 fields; ***p<0.001). Error bars show SEM. (B) Representative confocal fields of HEK293T cells stained with LipidTox^TM^ Red. (C) Representative graph and quantification of A375 cells undergoing a Mito Stress Test. Knockout of *NDST1* significantly increases the Baseline and Maximal oxygen consumption rate (OCR) of A375 cells (t-test, n=30 (10 wells x 3-time points) per genotype, *p<0.05, **p<0.01, ***p<0.001). The experiment was conducted in triplicate. Box and whisker plot shown in right panel.

HEK293T cells with integrated lentiviral vectors targeting *EXT1* or *NDST1* were used to evaluate the level of intracellular lipid when heparan sulfate elongation or sulfation was reduced (Fig. 2B). LipidTOX^TM^ staining was significantly lowered upon knockdown of either of these two heparan sulfate biosynthetic enzyme encoding genes. This is notable as a change in the pixel intensity distribution toward a greater representation of low-intensity pixels upon *EXT1* or *NDST1* knockdown compared to a scramble sequence lentiviral vector (Supplemental Fig. 1).

The results described above document morphological changes for markers of mitochondria and neutral lipids stored in intracellular lipid droplets upon reduction of key heparan sulfate biosynthetic enzyme encoding genes. To assess the functional consequences of changes in heparan sulfate levels or structure on mitochondria, oxygen consumption rates were measured with a Seahorse analyzer and a mitochondrial stress test. This test provides a measure of both baseline and maximal respiratory capacity. Significant increases in both baseline and maximal respiration rates were observed for A375 *Ndst1 -/-* cells compared to wild type A375 (Fig. 2C), and Hep3B *Ext1 -/-* compared to wild type Hep3B (Supplemental Fig. 2). These findings demonstrate that heparan sulfate modified proteins have a functional impact on the activity of mitochondria, with loss or partial loss of heparan sulfate biosynthesis increasing mitochondrial respiratory capacity.

### Knockdown of Ndst1 in mouse astrocytes reduces intracellular lipid levels

To evaluate the capacity of heparan sulfate to modulate attributes in cells affected by AD pathology, we examined intracellular lipids in wild-type mouse astrocytes as well as those expressing humanized *APOE3* and *APOE4* variants. Previous work has shown that astrocytes provide important metabolic support for neurons, and APOE variants affect this important capacity^43^. Astrocytes expressing the APOE4 variant showed elevated accumulation of intracellular lipid compared to wild-type or APOE3-expressing cells. Knockdown of *Ndst1* (to 60% of wild-type levels, measured by qPCR) using a lentiviral vector reduced intracellular lipid significantly in wild-type astrocytes, representing a greater than 4 fold reduction in lipid droplet volume (Fig. 3A). This was accompanied by significant reductions in *Perilipin 2* mRNA, the gene encoding a protein localized to the phospholipid shell of lipid droplets, compared to astrocytes transfected with a lentivirus bearing a scrambled sequence. Lentivirus-directed knockdown of *Ndst1* in astrocytes from mice expressing humanized *APOE3* and *APOE4* variants both display comparable and significant reductions in intracellular lipid compared to controls expressing a scrambled sequence lentivirus (Fig. 3B).

**Figure 3.**
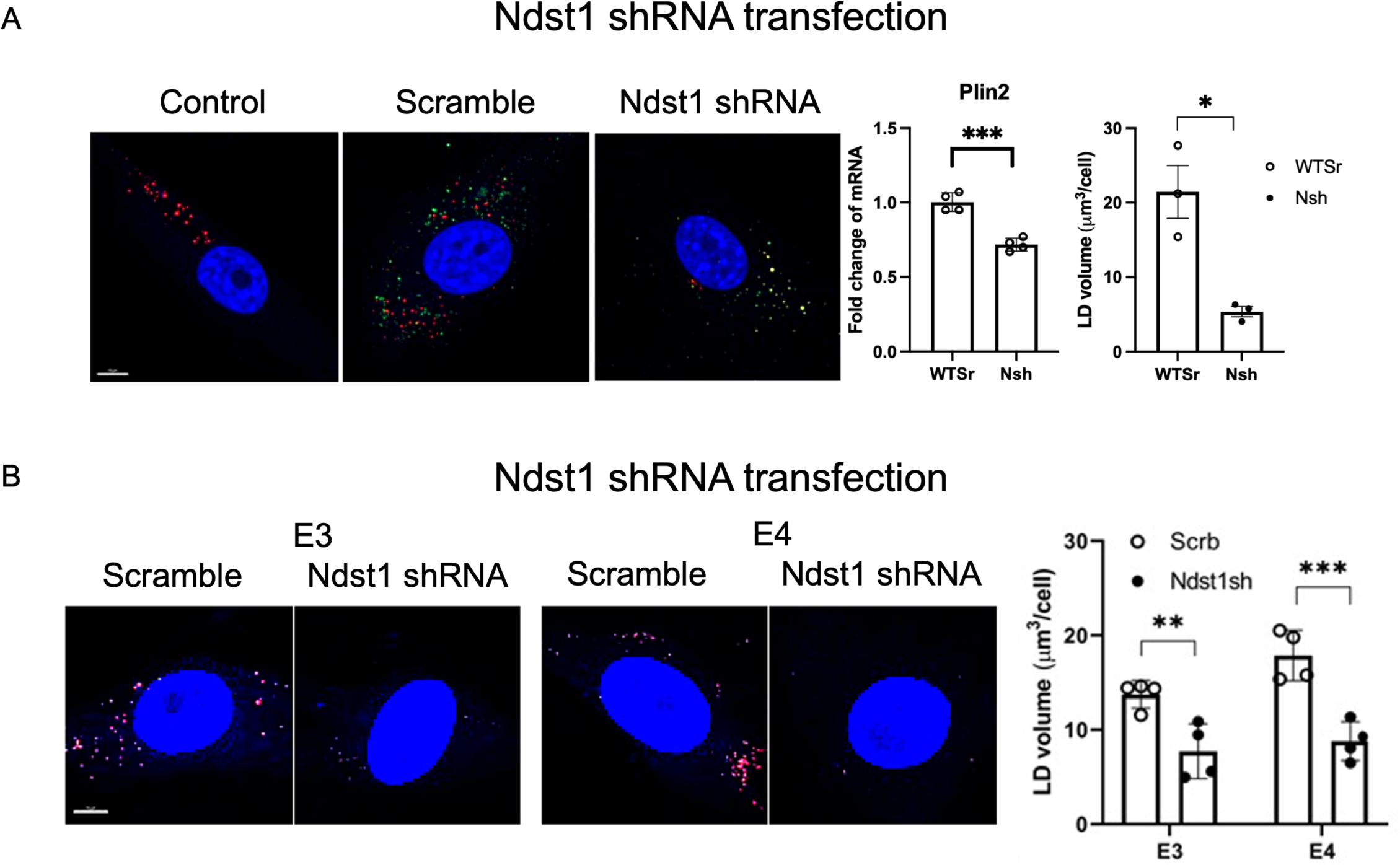
Knockdown of *Ndst1* in primary mouse astrocytes lowers levels of intracellular liposomes. (A) Micrographs showing levels of lipid droplets (LD), a marker for lentiviral transfection (green), and nuclear stain (DAPI, blue) in control, cells transfected with either a scramble sequence shRNA vector, or one targeting *Ndst1. Ndst1 shRNA* produced reduced levels of LD number and volume (panels on the right), as well as decreases in *Plin2* expression measured by qPCR. (B) The effect of *Ndst1 shRNA* was evaluated on astrocytes from animals expressing humanized variants of *APOE3* and *APOE4.* LD volumes were significantly reduced by *Ndst1* knockdown compared to scramble controls in astrocytes from either of these two genotypes [images show individual cells, nuclei (blue), and LD (red), and graphs provide quantitation of LD volumes. *p<0.05, **p<0.01, ***p<0.001). Error bars represent SEM.

### Presenilin/Psn deficit in Drosophila neurons produces degeneration suppressed by reducing heparan sulfate modification

Presenilin is one component of an enzyme complex, γ-secretase, a protease with a number of important substrates, including Amyloid Precursor Protein (APP) and Notch^44,45^. Mutations affecting *presenilin-1 (PS1)* function are responsible for a familial, early-onset form of AD. While *PS1* mutations show dominant inheritance, these variants are loss or partial loss-of-function, established by the demonstration that approximately 90% of 138 mutant proteins derived from human disease-associated variants and reconstituted into an active enzyme complex showed reduced Aβ peptide production^10,46^. These studies demonstrate that *PS1* partial loss-of-function models are relevant to understanding AD mechanisms. In vertebrate systems, compromising presenilin function results in mitochondrial dysfunction, accumulation of intracellular lipid, and disruption of autophagosome to lysosome trafficking^47^. With these observations in mind, the cellular and molecular phenotypes of partial loss-of-function changes in *Drosophila* γ-secretase were examined. These experiments were made possible by the development of effective RNAi constructs directed against two essential subunits of the γ-secretase complex, Presenilin (Psn), the catalytic subunit, and Nicastrin (Nct), an evolutionarily conserved component proposed to affect substrate selection^48^. All four protein components of human γ-secretase are represented in *Drosophila* and these two homologous enzyme complexes share common substrates. RNAi knockdown of either *Psn* or *Nct* mRNAs in neurons produces neurodegeneration in the adult brain and retina and has effects even when functional changes are conditionally restricted to adult neurons^49^.

We began by assessing the phenotypic severity of three RNAi constructs, two different *Psn* and one *Nct-*directed UAS-RNAi vectors expressed in the brain using a neuron-specific *Gal4* driver, *elav-Gal4.* Animals were evaluated for a startle-induced negative geotaxis assay, a locomotor reactivity response where animals move upward after a gentle tapping to the bottom of a plastic tube^50^. These findings confirmed previously published phenotypes, with *shPsn2* showing more severe behavioral deficits and *shPsn3* demonstrating a modest but significant deficit in behavior when compared to controls. Expression of *Nct* RNAi in neurons when animals were reared at 25°C was nearly completely lethal. The majority of the analyses described below therefore focused on *shPsn3*, the RNAi construct with an intermediate phenotype.

Earlier work has shown the capacity of reduced heparan sulfate biosynthesis to suppress cell degeneration, restore dysmorphic mitochondria, and reduce accumulation of ubiquitin-modified proteins in *parkin* mutants, a *Drosophila* model of *PARK2-*mediated Parkinson’s disease^20^. Partial reductions in the function of either of two genes involved in heparan sulfate biosynthesis, *sulfateless (sfl)*, encoding a homolog of *N-*deacetylase *N-*sulfotransferase (NDST1), or *tout velu (ttv)*, a glycosyl transferase responsible for heparan sulfate chain elongation (homolog of human *EXT1*), had dramatic effects on *parkin* mutants, restoring muscle cell morphology and flight activity^20^. We, therefore, examined the capacity of reduced heparan sulfate biosynthesis to affect neurodegeneration in the brain of *shPsn3* RNAi-expressing animals. Adult flies expressing *shPsn3* RNAi under the control of *elav-Gal4* were aged for 7 or 30 days prior to phalloidin and Hoechst staining of adult heads to visualize actin and nuclei in whole brain preparations^51^. Serial confocal sections of the entire brain were taken to identify vacuoles, areas of acellularity produced by neuronal loss (Figure 4A).

**Figure 4.**
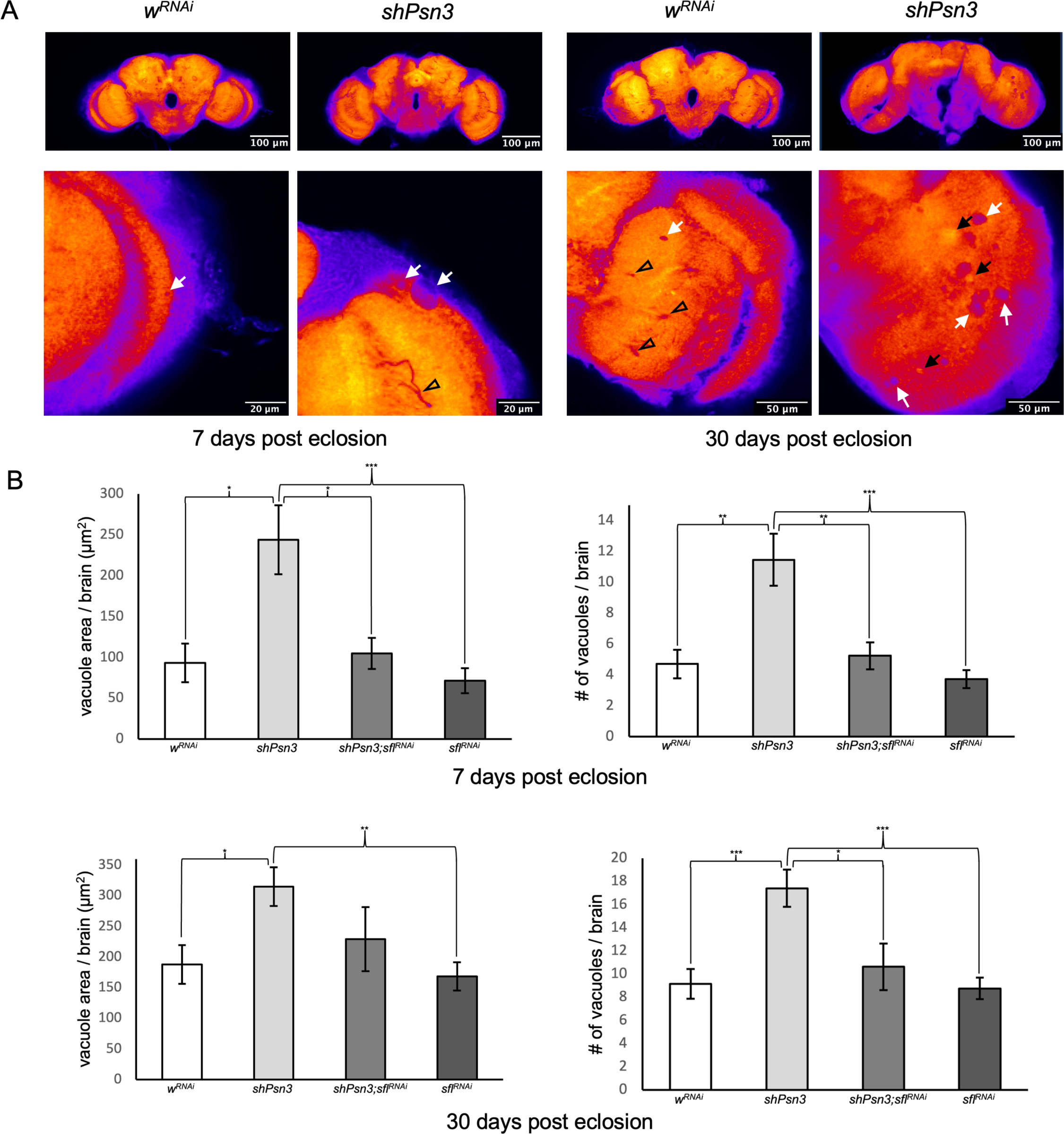
Reduction of *sfl* function reduces *Psn* knockdown-induced neurodegenerative vacuole formation. (A) Representative images of *Drosophila* brains stained with Alexa Fluor^TM^ 594-Phalloidin and imaged by confocal microscopy. Top row panels show low magnification images of the entire brain, bottom row, higher magnification (see scale bars). White arrows indicate neurodegenerative vacuoles. Black arrows indicate flares showing the higher staining intensity of phalloidin-associated with and near vacuoles. Black arrowheads indicate tracheoles, air conducting tubules, discernable in the serial confocal sections. (B) Bar graphs showing average vacuole area and count per brain in female *Drosophila*. Vacuole area and number of vacuoles per brain are significantly altered 7 days post eclosion (ANOVA Tukey test: *w^RNAi^* n = 10, *shPsn3* n = 24, *shPsn3;sfl^RNAi^* n = 14, *sfl^RNAi^* = 20; *p<0.05, **p<0.01, ***p<0.001), error bars represent standard error of the mean (SEM). Vacuole area and number of vacuoles per brain are significantly increased in *shPsn3* animals 30 days post eclosion and double *shPsn3;sfl^RNAi^* animals showed a significant reduction in vacuole number compared to *shPsn3 .* Vacuole area was reduced in these animals but did not achieve significance (ANOVA Tukey test: *w^RNAi^* n = 13, *shPsn3* n = 20, *shPsn3;sfl^RNAi^* n = 8, *sfl^RNAi^* = 20; *p<0.05, **p<0.01, ***p<0.001).

Consistent with earlier published work, expression of *shPsn3* in the adult brain produced a marked increase in neurodegeneration measured as the area of vacuolization or number of vacuoles per brain compared to control animals expressing a *w^RNAi^* construct (the *white* gene governs eye pigmentation) (Figure 4B). Knockdown of *sfl* transcripts with a *sfl^RNAi^* construct showed no difference from control animals with respect to levels of degenerative vacuoles. However, RNAi of *sfl* significantly reduced the level of degeneration in the brain mediated by *shPsn3* knockdown in animals aged 7 or 30 days (Fig. 4B, compare *shPsn3* to *shPsn3*; *sfl^RNAi^*). These findings demonstrate that *Psn* neurodegenerative phenotypes can be suppressed by modest reductions in *sfl* function and hence HSPG-mediated activities.

### Cellular processes regulated by Psn and heparan sulfate-modified proteins

The capacity of reduced *sfl* function to rescue neurodegeneration and behavioral deficits mediated by *presenilin* deficits prompted an analysis of cellular phenotypes with a focus on mitochondria, intracellular lipid, and autophagosome-related structures, all features known to be disrupted by *PS1* mutations in humans^47,52^. Given the documented effects of *PS1* mutations on many cell types and their involvement in conserved cellular functions, we examined the cellular phenotypes of *presenilin* and *sfl* knockdown in the larval fat body. The fat body serves as the principal metabolic organ in *Drosophila* where lipids and glycogen are stored and distributed to other tissues. Fat body cells are large (∼60μm) polyploid cells where intracellular organelles can be visualized in detail with specific fluorescent markers using serial confocal microscopy. Genes affecting mitochondria and neurodegeneration, including *parkin* and *pink1*, have been identified using RNAi screens and visualization of mitochondria in fat body cells^53^, demonstrating the utility of this preparation for assessing cellular phenotypes relevant to neural function and disease. Selective knockdown of genes in fat body cells is possible using UAS-RNAi-producing transcripts expressed under the transcriptional control of *r4Gal4*, a fat body-specific *Gal4* transgene. *r4Gal4* produces a reduction of *Psn*, and *sfl* mRNA by 32% and 28% of levels measured in control animals (*w^RNAi^*) respectively, determined by RNA Seq (see below).

### Psn and sfl knockdown in Drosophila fat body have opposing effects on mitochondrial structure and liposome organization

In order to evaluate the effect of presenilin function on mitochondria and intracellular lipids, *Psn-*RNAi expression was directed to the fat body. Mitochondria and intracellular lipids were visualized using the fluorescent probes MitoTracker^TM^ Red, and LipidTOX^TM^ by serial confocal microscopy, respectively. *Psn* knockdown had profound effects on the level and morphology of MitoTracker^TM^ labeled organelles in fat body cells (Figure 5). MitoTracker^TM^ signal intensity was significantly reduced in *shPsn3* preparations compared to control animals expressing *w^RNAi^.* The size of MitoTracker^TM^ labeled organelles was also affected by *Psn* knockdown (Fig. 5B), indicating that mitochondria are smaller upon reductions of *Psn* function. The effect of *shPsn3* on mitochondrial morphology was confirmed by transmission electron microscopy (TEM) and serial block face scanning electron microscopy (SBFSEM), revealing small, ellipsoid mitochondria in animals with *shPsn3*, compared to larger, branched mitochondria in control preparations (RNAi of *w* gene) or animals with *sfl^RNAi^* (Fig. 5D).

**Figure 5.**
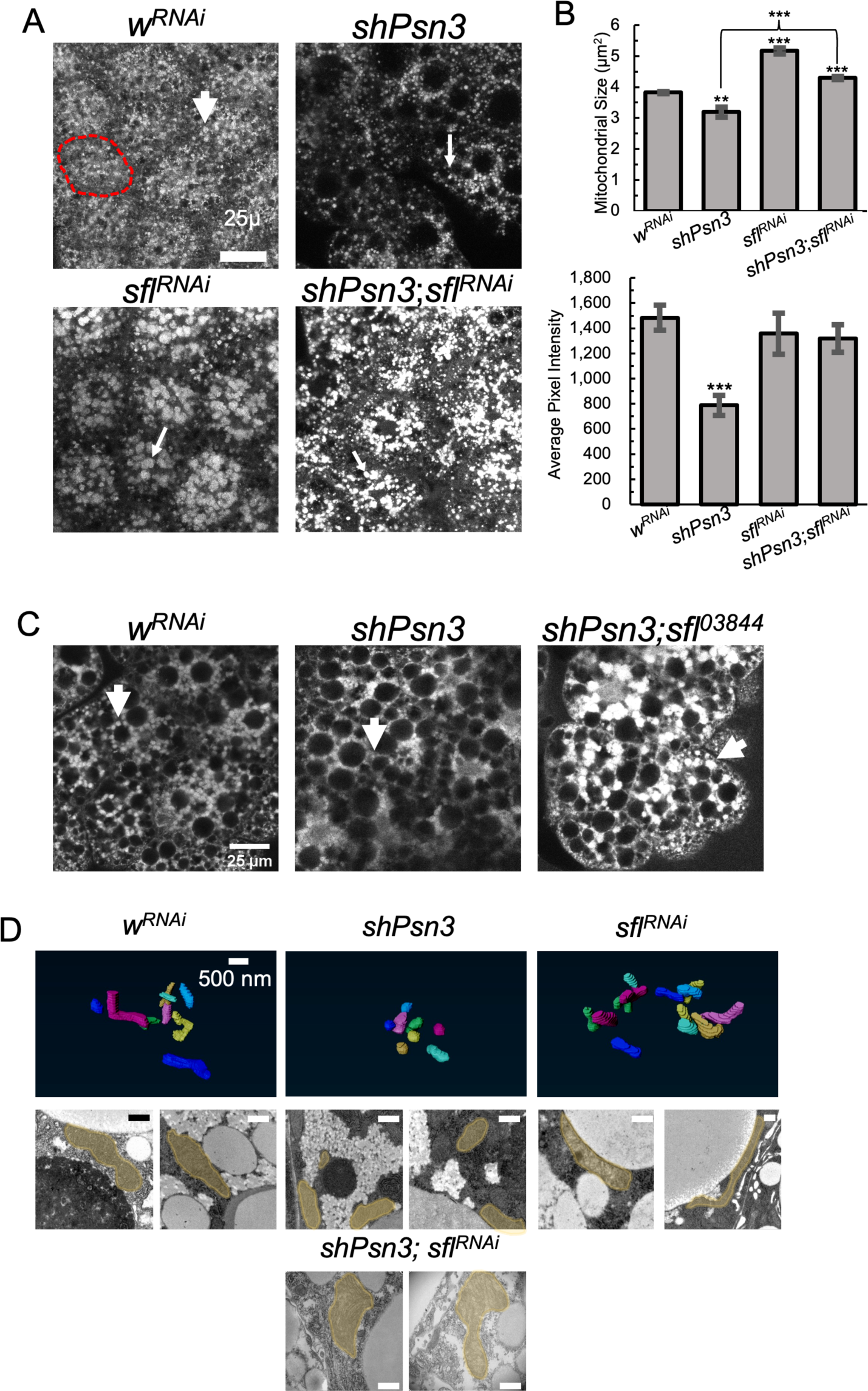
*psn* and *sfl* function alters mitochondrial number and morphology in fat body cells and *sfl* knockdown suppresses *psn^RNAi^-*mediated abnormalities. Mitochondria labeled with MitoTracker^TM^ Red and visualized by confocal serial section. (A) Images are projections of a stack of confocal images, showing the organization and morphology of mitochondria. The red dashed circle indicates the margins of one cell. White arrows mark individual mitochondria. Spheroid-shaped areas without MitoTracker^TM^ Red signal are the large liposomes found in fat body cells. (B) The measure of mitochondrial size shows *Psn* knockdown significantly reduces the size of MitoTracker^TM^ -positive structures compared to controls (*w^RNAi^*) whereas *sfl^RNAi^* has the opposite effect (increased size of MitoTracker^TM^ -positive structures), error bars represent SEM. (t-test p-values are in relation to *w^RNAi^* except where marked by bracket, **p<0.01, ***p<0.001). *sfl^RNAi^* rescues the mitochondrial phenotypes produced by *psn* knockdown, reflected both in morphology, and size of MitoTracker^TM^ -labelled structures. (p-values in relation to *w^RNAi^*, ***p<0.001). Significant differences were also measured for pixel intensity, with *shPsn3* showing reduced average pixel intensity compared to controls, and *sfl^RNAi^ shPsn3* animals restoring pixel intensity to control levels (*w^RNAi^)*. (C) Reducing *sfl* function with a characterized *P-element* mutant, *sfl*^03844^ also rescues mitochondrial number and morphology in fat body cells in the presence of *Psn* knockdown. Mitochondria labeled with MitoTracker^TM^ Red and shown in a single optical confocal section. (D). Top row panels show serial block-face EM assembly of mitochondria, providing a 3D rendering of mitochondrial morphology. Mitochondria in *shPsn3* are small spheroidal structures whereas both control (*w^RNAi^*) and *sfl^RNAi^* show more elongated mitochondria. The TEM images in the second and third rows of images show mitochondria (highlighted in translucent color) in these genotypes. Of note is the very elongated mitochondrion found in a *sfl^RNAi^* animal and the large, lobed mitochondrial found in *sfl^RNAi^ shPsn3* preparations.

We have previously shown that reductions in heparan sulfate biosynthetic enzyme encoding genes affect mitochondrial morphology in muscle, producing larger mitochondria via a process that requires an intact autophagy system^20^. We observe a similar phenotype with RNAi of *sfl* in fat body cells, with a more intense MitoTracker^TM^ signal and larger, labeled structures compared to *w^RNAi^* controls (Fig. 5A, B). SBFSEM and TEM confirmed the organization of mitochondria detected with MitoTracker^TM^ in *sfl^RNAi^* animals, with some large mitochondria evident in EM sections (Fig. 5D).

The morphology of liposomes was evaluated by confocal microscopy of neutral lipid fluorophore staining (LipidTOX^TM^) as well as TEM. Control animals (*w^RNAi^*) showed very regular liposomes, largely filling the entire cell (Fig. 6A panels). *shPsn3* animals had larger, spherical liposomes with an increase of irregularly shaped lipid-staining structures (Fig. 6A, red circled structures) compared to the control. TEM confirmed the presence of large and some fused liposomes in *shPsn3* animals (Fig. 6C). Also apparent were spherical structures smaller than liposomes, intermixed among them, and evident by their absence of staining.

**Figure 6.**
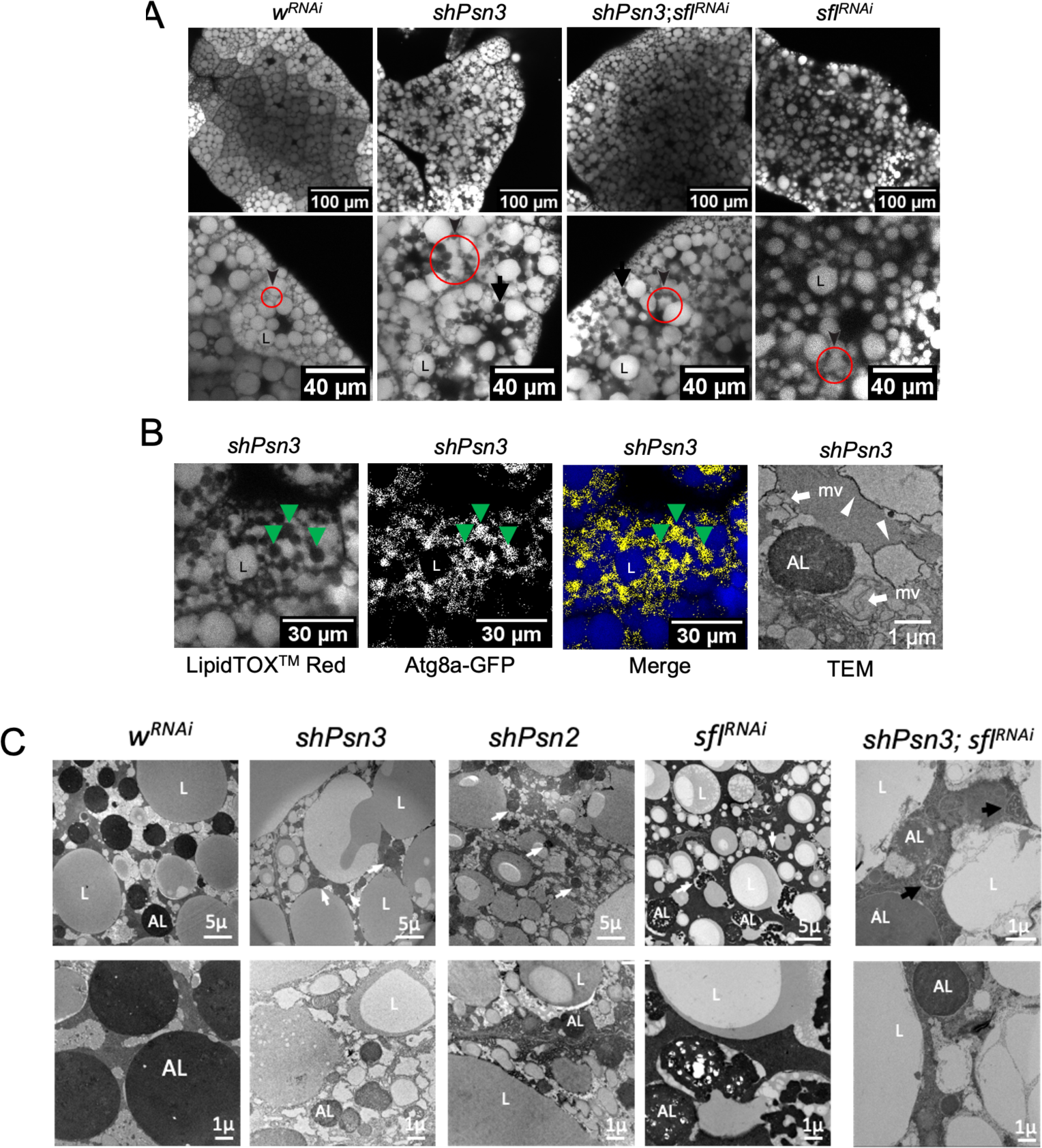
*sfl^RNAi^* partially reverts *psn* knockdown-induced alterations in liposome morphology. (A) Representative images of fat body stained with LipidTox^TM^ Red. Top row shows lower magnification image, while lower row shows higher magnification, more detailed view. Control fat body cells (*w^RNAi^*)show large, regular liposomes (L) filling the cell body. *shPsn3* results in changes in the spacing of liposomes and the appearance of irregularly shaped LipidTox staining structures (red circles and arrowheads). *sfl^RNAi^* produces a reduced density of liposomes, and also some irregularly shaped LipidTox staining structures. Arrows indicate vesicular-shaped gaps in the LipidTox^TM^ Red signal (see details in C). The letter ‘L’ labels liposomes. (B) Fat body cells stained with LipidTox^TM^ in animals expressing *Atg8a-GFP* transgene and *shPsn3*. The vesicular structures devoid of LipidTox staining show a GFP signal, indicating an accumulation of Atg8a (green arrowheads). The far-right panel shows a TEM micrograph of a *shPsn3* fat body cell with double membrane structures (white arrowheads), consistent with autophagosome morphology (relatively electron-lucent compared to autolysosomes (AL) as well as multi-vesicular bodies (arrows). These structures are found in fat body cells with compromised autophagy flux^54^. (C) TEM micrographs of fat body cells, showing the organization of autolysosomes (ALs) and liposomes (L). ALs appear as electron-dense structures, 2-5μ in control animals (*w^RNAi^*), but smaller and less electron-dense in animals with RNAi of *Psn.* Shown are TEM images from two independent *Psn* RNAi lines (*shPsn2, shPsn3*). Smaller ALs indicated by white arrows. In *sfl^RNAi^* animals, ALs display a different appearance, with electron-dense “cables” within a structure that can be seen fused with liposomes in some cases. Two micrographs of *shPsn3, sfl^RNAi^* preparations are shown (both higher magnification), with some restoration of AL size and electron density, as well as the appearance of ALs fusing with liposomes (black arrows) as seen in *sfl^RNAi^*.

These structures were not prevalent in control or *sfl^RNAi^* animals. To determine the nature of these intracellular organelles we examined the distribution of an autophagosome marker, GFP-Atg8a (Fig. 6B). Staining with both LipidTOX^TM^ and GFP-Atg8a showed these features to be Atg8a-positive and hence derived from an autophagosome-related organelle. The accumulation of these structures in *shPsn3* fat body cells indicates a change in autophagosome structure or traffic in animals with *Psn* knockdown. TEM sections from *shPsn3* animals also show an accumulation of double membrane organelles consistent with autophagosome morphology as well as multivesicular structures (Fig. 6B). These are cellular phenotypes characteristic of limited autophagosome maturation^54^ and further suggest that *Psn* knockdown compromises autophagy flux and an accumulation of Atg8a-positive structures. *sfl^RNAi^-expressing* animals also showed changes in liposome morphology, with a lower density of LipidTOX^TM^-staining structure compared to controls (Fig. 6A) and a greater representation of smaller liposomes. These findings were confirmed by TEM analysis (Fig. 6C).

*PSEN1* deficits are known to disrupt autophagosome traffic and this cellular abnormality is common to several neurodegenerative disorders. Conversely, reductions in heparan sulfate chain length or sulfation increase the trafficking of autophagosomes to the lysosome^19^. It was therefore of interest to examine autophagosome-related structures in fat body cells with *presenilin* or *sfl* knockdown. In TEM micrographs of wandering third instar larvae fat body cells, autolysosomes are prominent structures with a characteristic size of 2-5 μm, a single encompassing membrane, and containing varied electron-dense materials^54^. These are readily seen in TEM sections of control cells (Fig. 6C). In two different *Psn* RNAi-producing lines, TEM showed smaller, less electron-dense autolysosome structures, indicating that knockdown of *Psn* alters autophagy in these cells. *sfl^RNAi^* had a very different effect on autolysosomes, producing a membrane bound structure with a network of electron-dense “cables” on the interior. These autolysosomes were often seen in association with or having coalesced with liposomes (Fig 6C). These structures are consistent with the induction of autophagy and the liberation of lipids from liposomes in these animals^55^.

*Reduction of sfl function suppresses cellular abnormalities mediated by Psn knockdown* The cellular changes mediated by *Psn* or *sfl* knockdown were distinct, and in an apparently divergent manner for both mitochondrial and liposome morphology. Autophagy-related structures were also altered in different ways in *Psn* versus *sfl* RNAi-bearing animals. We examined these subcellular features in cells with reductions in both *Psn* and *sfl* and found a restoration toward wild-type morphology. Mitochondria were significantly larger and MitoTracker^TM^ signal levels were higher in the *shPsn3 sfl^RNAi^* double RNAi compared to *shPsn3* alone (Fig. 5B). Likewise, the distribution of liposomes was restored in the double knockdown toward the wild-type pattern (Fig. 6A). TEM supported the same conclusion with the appearance of nearly normal size autolysosomes along with those resembling those characteristic of *sfl* knockdown in the double *sh3Psn sfl^RNAi^* knockdown. This collection of phenotypes demonstrated the block or reversal of cellular deficits mediated by loss of *Psn* function by compromising the activity of heparan sulfate-modified proteins.

### Transcriptomics signatures of Psn, sfl, and Psn sfl double knockdown show that sfl knockdown restores wild-type patterns for key metabolic regulatory genes

The capacity of reducing heparan sulfate function to rescue cellular abnormalities of *presenilin* deficits prompted a search for molecular signatures that accompany that rescue. This strategy could identify genes that are functionally important for the pathology mediated by *presenilin* knockdown. Quadruplicate samples of RNA were isolated from the fat body of climbing third instar larvae of the following genotypes: *shPsn3, sfl^RNAi^, w^RNAi^* (controls), and *shPsn3; sfl^RNAi^* double knockdown. Genes with significantly changed expression levels in either *shPsn3* or *sfl^RNAi^* compared to *w^RNAi^* controls were identified, and those with opposing directions of change were determined (Fig. 7A). It is notable that the greatest number of genes with significant and opposite differential gene expression are down-regulated in *shPsn3* (Fig. 7B). For example, in the 200 genes with the greatest fold difference, 18 show opposite expression changes for the *shPsn3* down*, sfl^RNAi^* up gene set, and only one gene for *shPsn3* up, *sfl^RNAi^* down group. This suggests that the capacity of *sfl* knockdown to rescue *Psn* deficits lies largely with the correction of *shPsn3* down-regulated genes.

**Figure 7.**
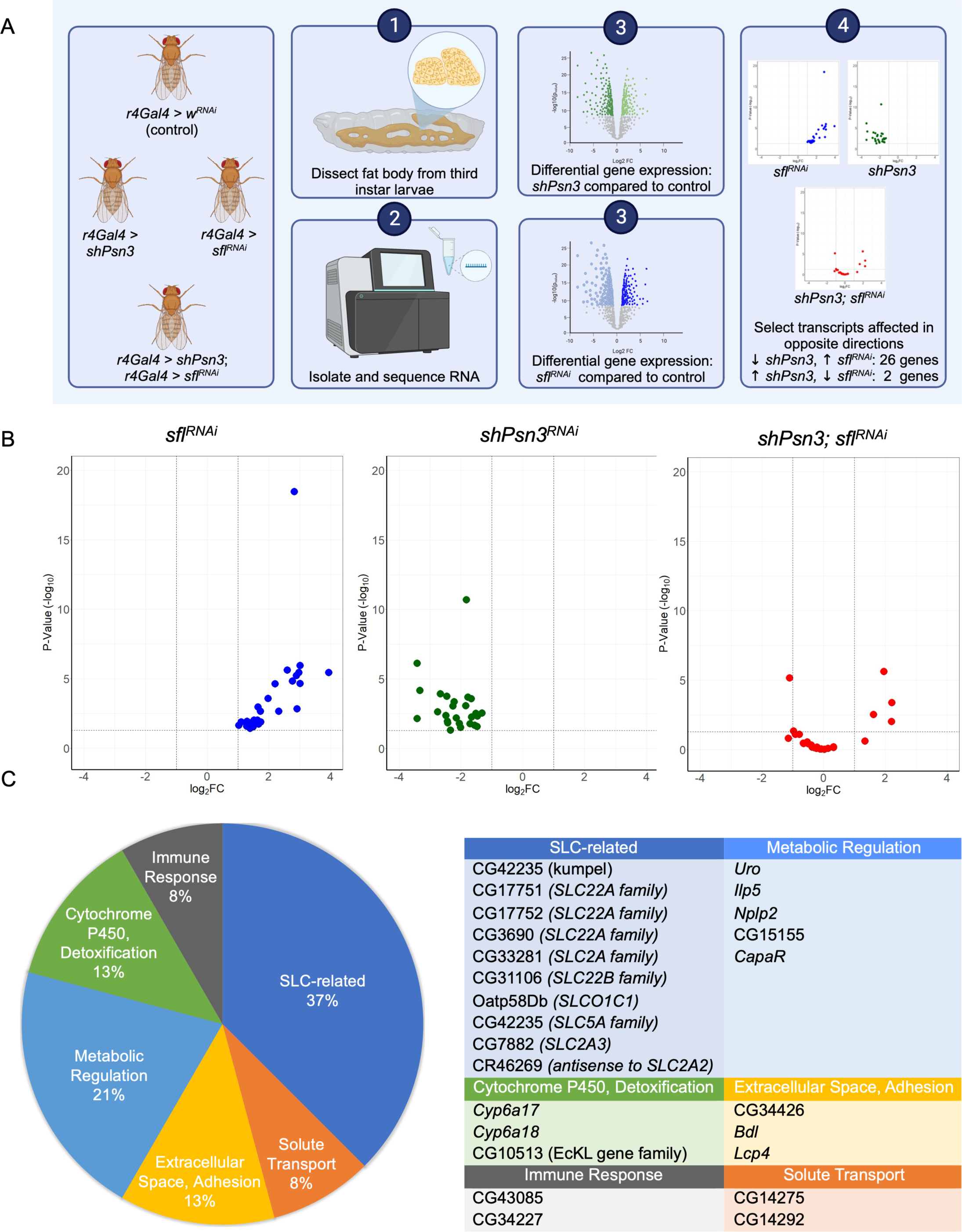
RNA seq analysis of fat body transcriptome shows *sfl^RNAi^-*mediated reversal of a set of differentially expressed genes from *shPsn3* animals. (A) Experimental scheme for identifying differentially expressed genes with opposing changes in *shPsn3* versus *sfl^RNAi^* and their expression in double *shPsn3; sfl^RNAi^.* (B) Volcano plots of genes significantly downregulated in *shPsn3* and upregulated in *sfl^RNAi^* (Y-axis: probability of expression changes, X-axis: fold change (FC) as log_2_. The far-right panel shows the volcano plot for those genes in the *shPsn3; sfl^RNAi^.* animals, demonstrating restoration of all but one gene to levels of control animals, or higher. (C) A breakdown of what genes show rescue of gene expression levels by functional class and listing of the affected transcripts, with their human homologs.

26 genes show this pattern of differential gene expression, significantly downregulated in *shPsn3* animals and upregulated in *sfl^RNAi^.* The expression of this set of genes was normalized in the *shPsn; sfl* double knockdown (Fig. 7B). Gene set analysis of these genes using g:profiler or PANGEA revealed significant enrichment after correction for multiple testing in GO biological processes Transmembrane transport and Reactome SLC mediated transporter (R-DME-425407, corrected p-value = 9.079e-4). A sizable proportion (37%) of these genes whose expression is decreased with *Psn* knockdown and displayed corrected expression mediated by *sfl* knockdown are members of the solute carrier gene family (Fig. 4C). CG42235 (*kumpel*) encodes a homolog of the SLC5A family of transporters, Na+-dependent monocarboxylate co-transporters. This family of SLCs in *Drosophila* includes *kumpel, rumpel*, and *bumpel*, and the latter has been directly demonstrated to transport both pyruvate and lactate^56,57^. Overexpression of *bumpel*, a gene expressed in largely the same glial cell set as *kumpel*, rescues autophagy defects and neuronal death in a *Drosophila* model of *C9orf72* neurodegeneration^56^. Two members of the *SLC2A* family of hexose transporters are also represented in the gene set rescued by *sfl*, CG33281, and CG7882. Three members of the *SLC22A* family were also downregulated in *Psn* knockdown animals and restored to normal levels of expression with *sfl^RNAi^.* This group of carriers transports organic ions, including carnitine, a molecule critical for fatty acid transport and oxidation in mitochondria^58,59^. Another group of transcripts suppressed upon *Psn* knockdown and restored to wild-type levels by simultaneous reduction of *sfl* expression affect metabolic regulation broadly and includes *Neuropeptide-like precursor 2 (Nplp2), Insulin-like peptide 5 (Ilp5)* and *Urate oxidase (Uro).* There are genetic interactions between insulin-like pathway signaling and uric acid accumulation in *Uro* mutants, suggesting the gene expression changes in *Ilp5* and *Uro* may be functionally connected^60^. *Nplp2* has recently been shown to affect lipid transport and homeostasis, serving as an exchangeable lipoprotein in the hemolymph^61^. *Ilp5* encodes a member of the insulin-peptide family and affects a number of metabolic processes^62^. These results collectively show that modest reductions in *Psn* function have dramatic effects on genes affecting metabolic processes that can be countered by modulation of HSPG function.

## DISCUSSION

A number of findings over the years have implicated heparan sulfate modified proteins in AD pathogenesis^15,63^. Recently, a dramatic example was reported for a member of a large kindred with a dominant disease-causing mutation in *PSEN1*^11^. This individual was cognitively intact well beyond the typical early onset of symptoms and was remarkable for being homozygous for a rare variant of *APOE3, APOE3 Christchurch.* This form of *APOE3* carries a mutation in the heparan sulfate binding domain, a change that essentially abrogated binding to heparin, a highly sulfated form of heparan sulfate. It was also reported in this study, that heparin-binding strength for other disease associated *APOE* variants was directly proportional to disease susceptibility (*APOE4>APOE3>APOE2>>ApoE3ch*). Recent work with mice bearing a humanized *APOE3ch* allele has provided evidence that the findings from that single individual are broadly applicable and point to a critical role for heparan sulfate function in *PSEN1*-mediated pathogenesis. The APOE3ch protein expressed in mice showed profound effects on microglial reactivity and Aβ-induced Tau seeding and spread^12^. Thus, the involvement of heparan sulfate modified proteins in PSEN1 and APOE function as well as AD pathogenesis seems evident. In this context it is important to bear in mind the diverse and important functions served by heparan sulfate modified proteins. HSPGs serve to assemble protein complexes to modulate a number of cellular processes, including endo- and exocytosis as well as signaling at the cell surface, providing co-receptor function for a myriad of protein growth factors. How then do HSPGs participate and influence AD pathogenesis?

The studies reported here begin with exploring the relationship between heparan sulfate modified protein function, and cellular and molecular signatures of AD pathogenesis in human cell lines and mouse astrocytes. We find that compromising heparan sulfate synthesis or modification has profound effects on autophagy, mitochondrial function and lipid metabolism in these vertebrate systems. The changes in these cellular functions were reflected in heparan sulfate-dependent changes in the transcriptome, including genes affecting cholesterol synthesis, lipid transport, autophagy, and mitochondrial biogenesis. There was also a remarkable enrichment of AD-associated genes identified by multiple GWAS studies in differentially expressed genes affected by loss of *EXT1*, a gene encoding a heparan sulfate polymerase. The cellular and molecular mechanism for HSPG modulation of AD-related changes was examined using a *Drosophila* model of *presenilin* mediated pathology. The results from this analysis were clear, reducing heparan sulfate function increased autophagy, mitochondrial function and biogenesis, and lowered intracellular lipid droplets, as observed in human cells and mouse astrocytes. Furthermore, reducing heparan sulfate modification by partial loss-of-function of *sfl*, an *NDST1* homolog in *Drosophila*, rescued cellular and molecular abnormalities in *Psn* defective animals, as well as the age-dependent neurodegeneration produced with partial reduction of *Psn* function.

The extensive genetic analysis of AD using GWAS, as well as whole exome and genome sequencing technologies, has made remarkable progress in identifying genes that confer susceptibility or causation of this disease. Collectively, these findings have identified shared genes and pathways affecting both familial, early onset as well as late-onset AD^4,5^. The current challenge is to identify the cellular and molecular events that are not only associated with disease but serve a functionally critical role in pathogenesis. Model organisms provide the potential for addressing this challenge by virtue of the genetic tools that can identify genes that modify or suppress disease phenotypes. The conservation of the genes and their functions shown to be critical in AD in the fruit fly *Drosophila* provides an opportunity to identify functionally critical events in the molecular and cellular pathogenesis. This model system also allows for the study of interactions suggested by human genetic findings that have important implications for identifying potential targets for pharmaceutical intervention.

### Transcriptome patterns and cellular events affected by heparan sulfate biosynthesis

Evaluating heparan sulfate-dependent transcription patterns using Hep3B cells bearing an *EXT1* knockout was informative with regard to the pathways and cellular processes governed by HSPG-mediated signaling. First, it provided molecular evidence for the profound effects of HSPG function on cholesterol biosynthesis and lipid transport. These findings link the effects of reducing HS synthesis on intracellular lipids stored in liposomes that we have documented for a variety of cell types, including mouse astrocytes. We also observed effects on transcripts affecting mitochondrial biogenesis and autophagy, all processes involved in neurodegenerative disease. Finally, there was an enrichment of genes with differential expression affected by *EXT1* among AD-associated genes identified by extensive GWAS study. HSPG-controlled transcripts overlap to a remarkable degree with this set of genes implicated in the development and susceptibility to late-onset AD.

Functional analysis of several human cell lines and mouse astrocytes demonstrated that HSPGs serve critical functions in autophagy, mitochondrial physiology, and lipid metabolism. Modest changes in HS structure can produce marked changes in mitochondria, intracellular lipid and autophagy flux. The changes affected by reducing heparan sulfate function counter the pathological shifts seen in AD and other neurodegenerative diseases, namely increases in autophagy, increased number and function of mitochondria, and lowered levels of intracellular lipid droplets.

We have used *Drosophila* to determine the functional interaction between *presenilin* and HSPGs as one model for evaluating the impact of HSPGs on neurodegeneration. In particular, the cellular, and molecular phenotypes of reducing the function of the *Drosophila presenilin* gene, *Psn*, and a gene critical for heparan sulfate biosynthesis, *sfl* were examined. Heparan sulfate-modified proteins are known to regulate processes disrupted by *presenilin* deficits, autophagy in particular. In both *Drosophila*^19,20^ and mice^64,65^, compromising the function of either heparan sulfate biosynthetic enzyme-encoding genes or individual heparan sulfate-modified core proteins produces an increase in autophagy flux. For example, reductions in Perlecan (Hspg2) in mice has profound effects on autophagy^64^, as well as mitochondria and lipid metabolism^65^, events linked to changes in autophagy. In human hepatocellular carcinoma, elevated *Glypican-3* expression is associated with the suppression of autophagy, and reducing *GPC3* elevates autophagy levels and suppresses cell growth^66–68^. In contrast to the effects of lowering heparan sulfate-modified protein function, mutations in *PSEN1* compromise autophagosome to lysosome flux^69–72^, and activating autophagy can restore autophagy levels and rescue mitochondrial abnormalities^73^. These findings point to the regulation of autophagy as a key element of AD pathogenesis, and heparan sulfate-modified proteins as an important determinant of autophagy flux.

*Psn and sfl knockdown have opposing effects on key cellular processes, and modest reductions of sfl suppress Psn-mediated cell deficits and neurodegeneration* Previous work established that reductions in *presenilin/Psn* or *Nicastrin*, another component of the γ-secretase complex conserved in *Drosophila*, in adult neurons produce age-dependent degeneration^49^. We used this system to explore not only the interaction of *Psn* with heparan sulfate biosynthesis in neurodegeneration but evaluate the cellular and molecular signatures in fat body cells, the principal metabolic storage and transport organ in *Drosophila. Psn* and *sfl* knockdown showed divergent effects on mitochondrial number and morphology, as well as intracellular lipid and autolysosome structure seen with EM and confocal visualization of fluorescent-tagged markers. Consistent with these different effects on cellular features, *sfl* knockdown was able to suppress both neurodegeneration and the spectrum of cellular phenotypes produced by partial loss-of-function *Psn* knockdown. It is notable that modest changes in the function of either of these genes produced readily discernable phenotypes. RNA seq analysis of fat body cells with knockdown achieved with RNAi vectors showed 30-40% reductions in mRNA and clear effects on cell phenotypes and cell loss in the nervous system.

Knockdown of *Psn* in fat body cells produced differential gene expression changes, as did *sfl^RNAi^.* Comparing the differential expression patterns showed a subset of genes with opposite directions of change. The vast majority of these genes were downregulated in *Psn* knockdown animals and rescued to normal levels with simultaneous reduction of *sfl* mRNA levels. The identity of these genes, reflecting a molecular rescue signature, showed a large representation of solute carrier transporters, molecules inherently affecting metabolism and energy balance. One SLC family member was of particular interest, CG42235 (*kumpel*), encoding a homolog of the SLC5A family of transporters, Na^+^-dependent monocarboxylate co-transporters. *Kumpel* and another member of this family, *bumpel*, have been shown to transport lactate and pyruvate^57^, and when overexpressed, can rescue neuron loss mediated by expression of *C9orf72*, a human disease gene affecting frontotemporal dementia and amyotrophic lateral sclerosis^56,74^. Monocarboxylate and lipid transporters have also been shown to be vital in neuron-glia interaction in responding to ROS damage affected by mitochondrial dysfunction in neurons^75^. One gene affected by *Psn* knockdown and rescued by compromising HS sulfation is *Nplp2*, a lipid transport protein that responds to thermal stress, promoting survival^61^. *Nplp2* has functions that parallel *APOE* in vertebrate systems, namely serving as an exchangeable lipoprotein that can be soluble in hemolymph or blood and also associate with VLDLs^61^. Our transcriptomic analysis of *Psn* rescue by *sfl* has identified classes and gene family members shown to be critical in two other models of neurodegeneration in *Drosophila.* These findings emphasize the participation of HSPGs in cellular signaling systems affecting neuronal survival and point to processes critical in neurodegeneration, namely monocarboxylate and lipid transport.

### HSPGs and AD pathogenesis

It has been known for some time that HSPGs are components of AD-pathological structures, namely neuritic plaques containing APP-derived peptides. Compromising HS biosynthesis reduces the levels of these pathological structures in mouse models of AD as does ectopic expression of HS degradative enzymes^76,77^. It is also known that HSPGs affect the internalization and propagation of Tau aggregates, providing structures that seed intracellular fibrils^78^. These studies all suggest mechanisms where HSPGs contribute to AD pathogenesis and therefore point to HS synthesis as a potential therapeutic target. The studies here provide another dimension; HSPGs affect signaling events that influence cellular processes known to be early and important for the development of neurodegenerative disease. Our studies show that HSPGs regulate autophagy, mitochondrial structure and function, and lipid metabolism, all important processes in neurodegeneration. We have provided direct evidence for HSPGs affecting neurodegeneration and cellular deficits mediated by reductions in *presenilin* function. HSPGs are cell surface signaling proteins, and their effect on *presenilin-*mediated cell defects indicates that changes in signaling are an important contributor to AD pathogenesis. It is intriguing that GSEA pathways with suppressed signaling in *EXT1 -/-* cells are linked to aberrant signaling (Supplemental Table 4: Signaling by PDGFR in Disease and Signaling by BRAF and RAF1 Fusions). Perhaps reducing heparan sulfate biosynthesis suppresses multiple signaling processes that promote AD pathogenesis.

## AUTHOR CONTRIBUTIONS

S.B.S., conceived the experiments in *Drosophila*, and human cell systems, obtained the funding and lead the writing and editing of the manuscript. R.W. designed and conducted the FACS analysis and edited the manuscript. F.Y. and W.W. designed and conducted the mouse astrocyte experiments, contributed the corresponding figures and edited the manuscript.

N.S. conducted and designed both *Drosophila* and human cell line experiments, assisted in manuscript assembly and editing, and guided the efforts of A.C., A.K., R.B., R.M., S.S., M.O., and M.R. on both *Drosophila* and human cell line work. A.C. conducted the neurodegeneration experiments and assisted in manuscript assembly and editing.

## ACKNOWLEDGEMENTS

This work is supported by NIH/NIA grant (R21AG070843) and funding from the Penn State Eberly College of Science Lab Bench to Commercialization grant to S.B.S. We also want to thank Missy Hazen and the Huck Institutes Microscopy Core facility for assistance with EM studies as well as the. Huck Institutes’ Flow Cytometry Core Facility for use of the BD LSRFortessa Special Order Research Product (SORP) and Dr. M. Rajeswaran for assistance with sample preparation. We also want to thank Lindsey Swanson and Sophia DeGuara for contributions to some of the *Drosophila* confocal microscopy experiments, Grace O’Sullivan, and Jolyn Toyomura for human cell line culture experiments, Uzair Isaiah for *Drosophila* culturing and *Nct* and *Psn* experiments, and Peter Mason and Tuyen Pham for coding and graphics expertise. We thank Maitreya Das, Deepro Banerjee, and Istvan Albert for their recommendations and assistance with the transcriptome analysis. We are indebted to Santhosh Girirajan for many helpful discussions and careful reading of the manuscript. Our thanks are extended to Teresa Niccoli for sharing of unpublished work and her thoughtful recommendations on the manuscript.

## STAR Methods

### RESOURCE AVAILABILITY

#### Lead Contact

Further information and requests for resources should be directed to and will be fulfilled by the Lead Contact, Scott Selleck.

#### Materials Availability

Fly and cell lines used in this study are available upon request.

#### Data and Code Availability

Raw RNA Seq Data from this study are provided in Tables S2 and S3. This paper does not report any original code.

Any additional information required to reanalyze the data reported in this paper is available from the lead contact upon request.

##### EXPERIMENTAL MODEL AND STUDY PARTICIPANT DETAILS

All animal experiments were approved by the University of Arizona Institutional Animal Care and Use Committee (IACUC). Humanized ApoE3 (029018) and ApoE4 (027894) knockin mice (homozygous) and the wild-type C57BL/6J mice were obtained from the Jackson Laboratory and bred at the University of Arizona animal facility. All mice were housed in a temperature (23C) and humidity-controlled room with a 12-h light and 12-h dark cycle with ad libitum access to water and standard laboratory global 2019 diet (23% calories from protein, 22% calories from fat, 55% calories from carbohydrate, 3.3 kcal/g; Envigo) and cared in facilities operated by University Animal Care. 3-month male mice were used for primary astrocyte isolation and in vivo / ex vivo assays. Timed pregnant mice of designated genotype were prepared for the isolation of E17 embryonic hippocampal neurons.

Fly strains were raised on standard cornmeal/sucrose/agar media at 25°C under a 12 h day/12 h night cycle. w^1118^ served as the wild-type stock. [UAS*-Atg8a* RNAi (43097)] and [UAS-*w* RNAi (30033)] RNAi strains were obtained from the Vienna *Drosophila* Resource Center (VDRC). [UAS-*sfl* RNAi (34601)], [sfl^03844^ (5575)], and [c*155 elav-gal4; UAS-dcrII (25750)] were obtained from the Bloomington *Drosophila* Stock Center (BDSC). [UAS-shPsn2] and [UAS shPsn3] flies were a generous gift from J. Shen and Norbert Perrimon, Harvard Medical School, Boston, MA. [shPsn3; UAS-*sfl* RNAi] and [shPsn3RNAi; *sfl*^03844^] stocks were generated in lab.

A375, Hep3B, and HEK293T Cell lines were cultured in DMEM supplemented with 10% fetal bovine serum (FBS) and 1% penicillin/streptomycin at 37°C, 5% CO_2_ (Gibco).

### METHOD DETAILS

#### *Drosophila* brain dissection and assessment of neurodegeneration

Following eclosion, flies were sorted by sex and aged for 7 or 30 days. Whole flies were fixed in 4% PFA containing 0.5% Triton X-100 (0.5% PBS-T) for 3h with nutation at 20°C. The samples were washed 4 times with 0.5% PBS-T for 15 min with nutation at 20°C. Brains were dissected from whole flies in 0.008% PBS-T and placed in staining solution (DAPI at 1:1000 and Phalloidin at 1:100 in 0.5% PBS-T) for 16-24h with nutation at 4°C. Following staining, brains were washed 4 times with 0.5% PBS-T for 15 min with nutation at 20°C and once with 1xPBS for 30 min with nutation at 20°C to remove any residual detergent or stain. A SecureSeal^TM^ imaging spacer was applied to each microscope slide, and brains were mounted anterior side up with SlowFade^TM^ Gold Antifade Mountant (without DAPI). A more detailed version of this procedure can be found in^51^. Full brain samples were imaged non-sequentially on an Olympus FV3000 confocal microscope at a step size of 2 µm. Images were analyzed in Fiji ImageJ using the trace and measure tools. The Look Up table was set to Fire, and the maximum pixel intensity was adjusted to provide optimal visualization of vacuoles. Once a vacuole could be distinguished from a tracheole, it was traced in the slice in which the diameter was the largest. The total number of vacuoles per brain and the total area of vacuoles per brain were summed, respectively. ANOVA statistical analysis (one-way, Tukey comparison) was performed in Minitab to confirm significance.

#### Negative geotaxis assay

The behavioral abilities of Drosophila were measured via a startle-induced negative geotaxis protocol^50^. Upon eclosion, flies were separated by sex and aged for 3-6 days. For each geotaxis assay, 10 flies were placed into the apparatus and a startle response was induced by firmly tapping the apparatus on a mat until all flies had fallen to the bottom. The number of flies above the 8 cm line after 10 seconds was recorded and the flies were given a 60-second rest period before the next trial was performed. An average of 5 repetitions per vial of flies was taken as a single data point.

#### MitoTracker^TM^ Red staining of fat body

The fat body of third instar *Drosophila melanogaster* larvae were dissected in PBS and transferred to a 100 nM solution of MitoTracker^TM^ Red CMXROS. They were incubated at room temperature for thirty minutes out of light. The fat bodies were washed in PBS and then fixed in a 4% v/v PFA solution where they were incubated at room temperature for thirty minutes out of light. After fixation, the fat body was washed in PBS and transferred to the glass imaging slide. ProLong™ Gold Antifade Mountant with DNA Stain DAPI was applied to the fat bodies as a mounting medium with counterstain. The fat bodies were imaged with an Olympus FV3000 confocal microscope.

Images of fat body stained with MitoTracker^TM^ Red CMXRos were processed in Fiji ImageJ on a maximum intensity projection. An auto-threshold was applied to the images through Max Entropy. A watershed was applied and segmented mitochondria. Particles were selected for those between 1-10 µm in area and between 0.25-1.00 in circularity.

The size of mitochondria was analyzed using the built-in particle analysis tool in Fiji ImageJ on a maximum intensity projection. A 200x200 pixel box was used as a sample area, with the upper left pixel being the locator pixel that corresponded to the coordinates of the box. A random number generator was used to select two numbers between 0 and 824 to generate a random (x,y) coordinate to place the box. If the box was not entirely over a region where signal was present, the coordinates were reselected. Each sample was analyzed using this method three times, ensuring that the analyzed areas did not overlap.

#### LipidTox^TM^ Red staining of fat body

The fat body of third instar *Drosophila melanogaster* larvae were dissected in PBS and then fixed in a 4% v/v PFA solution where they were incubated at room temperature for thirty minutes out of light. After fixation, the fat bodies were washed in PBS. Fat bodies were then transferred to a LipidTox Red solution diluted 1:999 in buffer and incubated for thirty minutes at room temperature out of light. After staining, the fat bodies were transferred to a glass imaging slide. ProLong™ Gold Antifade Mountant with DNA Stain DAPI was applied to the fat bodies as a mounting medium with counterstain. The fat body tissue was imaged with an Olympus FV3000 confocal microscope.

#### Transmission electron microscopy

Fat body tissues were dissected out of third-instar larvae and stored overnight in a refrigerator in Modified Karnovsky’s fixative. Samples were washed three times in 0.1M cacodylate buffer and subsequently fixed for one hour in 1-2% Osmium tetroxide in 0.1M cacodylate buffer. Samples were washed two times with 0.1M cacodylate buffer and one time with milliQ water before undergoing an en bloc stain for one hour in filtered 2% aqueous Uranyl acetate. Samples were dehydrated in progressively increasing ethanol concentrations (50%, 70%, 85%, 90%, 100%, 3xEM grade ethanol) followed by three acetone dehydrations. Samples were infiltrated with Spurr’s resin and imaged.

Samples were examined using FEI Tecnai G2 Spirit BioTwin and FEI Talos 200C microscopes.

#### RNA Isolation

RNA was isolated according to the Macherey-Nagel NucleoSpin® RNA Kit protocol for cultured cells and tissue. Briefly, Hep3B or *Drosophila melanogaster* fatbody cells were lysed in 350 µl Buffer RA1 and 3.5 µl β-mercaptoethanol and then filtered through a NucleoSpin® Filter column through centrifugation for 1 minute at 11,000 x *g*. The lysate was then mixed with 350 µl 70% ethanol and passed through a NucleoSpin® RNA Column at 11,000 x *g* for 30 seconds in order to bind the RNA to the column. The binding step was repeated three additional times for lysate from fatbody. 350 µl of Membrane Desalting Buffer was added to the NucleoSpin® RNA Column and centrifuged for 1 minutes at 11,000 x *g*. 95 µl of a DNase reaction mixture was incubated on the column for 15 minutes at room temperature to digest DNA. After digestion, 200 µl of Buffer RAW2 was applied to the column and centrifuged for 30 seconds at 11,000 x *g* to inactivate the rDNase. 600 µl of Buffer RA3 was then added to the column and centrifuged for 30 seconds at 11,000 x *g*. 250 µl of Buffer RA3 was applied to the column and centrifuged for 2 minutes at 11,000 x *g* to dry the membrane. 60 µl of RNase-free H_2_O was applied to the column and centrifuged at 11,000 x *g* for 1 minute in order to elute the RNA. Hep3B cells were grown to 70% confluency in a six-well plate at 37°C and 5% CO_2_ in DMEM with 10% FBS and 1% Penicillin/Streptomycin.

#### RNA Sequencing

RNA sequencing was performed by Psomagen using the Illumina Truseq stranded mRNA library, a 40bp sequencing coverage, 40 million total reads, and a read length of 150bp PE on a NovaSeq6000 S4. All samples passed Psomagen Quality Control checks for both RNA and cDNA. RNAseq quality was validated by checking for a Q30 greater than or equal to 85% through NovaSeq 6000 and NovaSeq X Plus analysis, a Q30 greater than or equal to 80% through HiseqX analysis, and a Q30 greater than or equal to 70% for 2 x 300bp through MiSeq analysis. Low-quality bases with a Q less than 20 were trimmed. Subsequently, reads with more than 10% low-quality bases were trimmed. Adapter sequences were then trimmed from remaining reads. Ribosomal RNAs and RNAs mapped to incorrect genomes were filtered out of the data set. Alignment coverage, depth, and pair-end mapping information were generated based on the percentage of mapped reads, percentage of reads mapped onto specific genomic regions defined by GTF or GFF, 3’ and 5’ bias patterns, the properly paired read percentage, and the strandness of the alignment.

Transcriptomics analysis was performed in BasePair, in which one Principle Component covered 90.53% of variants in HEP3B data and in the mid-40 percentile for different genotypes in *Drosophila* data with quadruplicate samples.

#### RNA Sequencing data analysis

RNA sequencing data was analyzed, and Differential Expression data was generated by Basepair. Data was aligned to the appropriate genome using STAR and gene expression levels were quantified using featureCounts. Differential Expression data was generated using DESeq2 and pathway enrichment analysis was performed using GSEA.

#### MitoBlue staining of human cultured cells

Tissue culture cells were plated in DMEM with 10% FBS and 1% Penicillin/Streptomycin on Ibidi 12-well chamber slides at 30,000 cells/well the day before they were to be stained. The cells were kept overnight at 37°C and 5% CO_2_. On the day of the experiment, the complete DMEM was removed and a 5 µM MitoBlue solution in DMEM containing no FBS or PenStrep was applied to the cells. The cells were incubated for 30 minutes at 37°C out of light. The staining solution was aspirated, and the cells were washed with PBS. 4% v/v PFA was applied to the cells, which were then incubated for 30 minutes at room temperature out of light. The PFA and rubber chambers were then removed and Vectashield was applied as a mounting medium. The cells were imaged with an Olympus FV1000 microscope. Images of Hep3B cells stained with MitoBlue were processed in Fiji ImageJ on a maximum intensity projection. An auto-threshold was applied to the images through Max Entropy. A watershed was applied and segmented mitochondria. Particles were selected for those between 1-10 µm in area and between 0.25-1.00 in circularity.

#### LipidTOX^TM^ Red staining of human cultured cells

Tissue culture cells were plated in DMEM with 10% FBS and 1% Penicillin/Streptomycin on Ibidi 12-well chamber slides at 30,000 cells/well the day before they were to be stained. The cells were kept overnight at 37°C and 5% CO_2_. On the day of the experiment, the DMEM was removed and 4% v/v PFA solution, a fixative, was applied to the cells for 30 minutes at room temperature out of light. The PFA was aspirated, and the cells were washed with PBS. <u>LipidTOX^TM^</u> Red diluted 1:1000 in buffer was applied to the cells and they were incubated for 30 minutes at room temperature out of light. The <u>LipidTOX^TM^</u> Red solution and rubber chambers were then removed and Vectashield was applied as a mounting medium. The cells were imaged with an Olympus FV1000 microscope. Maximum projection confocal images of HEK293T cells were analyzed for differences in <u>LipidTOX^TM^</u> Red signal intensity. A 145x145 pixel box was used as a sample area, approximately ten cells were included in each pixel box. A histogram was collected of each sample area, with 256 bins collected in each histogram. The scale of pixel intensities was set from 0-4096. Each sample was analyzed using this method three times, ensuring that the analyzed areas did not overlap.

#### Preparation of fluorescently tagged guanidinylated neomycin-Cy5 conjugate and uptake experiments

Biotinylated guanidinylated neomycin (GNeo-biotin) was synthesized, as previously described^42^, and stored in water at -20 °C. After thawing at room temperature, GNeo-biotin was diluted into PBS to 2.5 µM. To this solution, streptavidin-Cy5 (ST-Cy5, Thermo Fisher) was added to achieve a final molar ratio of 1:5 of fluorophore to biotin. To ensure completion of the biotin-streptavidin reaction, the solution was gently mixed and allowed to incubate at room temperature, shielded from light, for 20 min. Following this incubation, the GNeo-ST-Cy5 conjugate was diluted to the desired concentration in DMEM media. A375 wildtype or NDST1-/- knockout cells were plated at 200,000 cells/well in 6-well tissue culture plates and let adhere overnight. The next day, the cells were washed with PBS and incubated with fluorescent-tagged GNeo-biotin (10 nM) for 1 hour at 37 °C under an atmosphere of 5% CO2. Control cells were treated with ST-Cy5 only. After incubation, cells were washed twice with PBS, released with 0.05% trypsin/EDTA (Corning), and analyzed by flow cytometry.

MitoTracker^TM^ Red and LipidTOX^TM^ Green uptake studies and flow cytometry A375 wildtype or NDST1-/- knockout cells were plated at 200,000 cells/well in 6-well tissue culture plates and let adhere overnight. For drug treatments, DMSO or rapamycin (10 µM) was added the next day, then cells were incubated for 24 hours at 37 °C under an atmosphere of 5% CO2. The next day, the cells were washed with PBS and incubated with MitoTracker^TM^ Red (1:1000, Thermo) or HCS LipidTOX<u>^TM^</u> Green (1:500, Thermo) diluted in DMEM media for 1 hour at 37 °C under an atmosphere of 5% CO2. After incubation, cells were washed twice with PBS, released with 0.05% trypsin/EDTA (Corning), and analyzed by flow cytometry. Flow cytometry experiments were performed using a CytoFLEX S (Beckman Coulter) flow cytometer (≥10,000 events/sample), and raw data were analyzed using FlowJo Analytical Software v10.8 (Becton Dickinson). Cells were gated according to forward and side scattering. The extent of uptake was quantified using the geometric mean of the fluorescence intensity versus control samples. These values were plotted and further analyzed using GraphPad Prism v10.0.

#### Seahorse Measurement of Oxygen Consumption Rate

The cell’s rate of oxidative metabolism of glucose was evaluated with XF Cell Mito-stress Assay in a Seahorse XFe24 analyzer which allows real-time metabolic analysis in live cells. Three cell lines, A375, Hep3B, and HEK293T, were plated on Agilent Seahorse XF24 Cell Culture Microplates (30,000 cells/per well) and were placed in a CO2 incubator overnight. The sensor cartridge used from an Agilent Seahorse XFe24 Extracellular Flux Assay kit was hydrated with Seahorse XF Calibrant (1000 μL/well) overnight in a non-CO2 incubator. Cells were subsequently washed with XF assay media [0.5 mM Glutamine (200 mM solution), 0.5 mM Glucose (1.0 M solution), 0.5 mM Pyruvate (100mM solution), 48.5 mM Seahorse XF DMEM assay medium, pH 7.4]. After washing, cells were incubated in respective media (37 °C, 1 h) in a non-CO2 incubator, allowing for temperature/pH equilibration before each measurement in the metabolic analyzer. Each plate contained four wells not seeded with cells to serve as blank controls for temperature-sensitive fluctuations. In the case of HEK293T, there were six control wells, to divide specified wells evenly between three genotypes. The lyophilized drugs from the Seahorse XF Mito Stress kit were diluted in Seahorse XF assay media and loaded into the corresponding injection ports in the sensor cartridge. Following measurements of resting respiration, cells were subsequently treated with Oligomycin (1.5 μM) for the assessment of non-phosphorylating oxygen consumption rate (OCR), with the mitochondrial uncoupler FCCP (1.0 μM) for the evaluation of maximal OCR, and with a combination of Rotenone and Antimycin A (both at 0.5 μM) for estimating extra-mitochondrial OCR. In all cases, three measurements were recorded, each over a 5-8 min interval including mixing and incubating. At the end of the Seahorse run, to allow comparison among different experiments, data were normalized to the total cell amount per well. In all cases, the metabolic parameters of the assay–basal and maximal respiration, proton leak, and ATP production through oxidative phosphorylation– were calculated with Agilent/Seahorse XF Report Generator software and expressed as OCR in pmol/min. The results are illustrated as a representative from at least three independent experiments performed in triplicate or means ± standard error from at least three independent experiments performed in triplicate.

#### Isolation and culture of Primary Astrocytes

Primary astrocytes were isolated as described in ref^43^. Fresh mouse brain tissue without cerebellum or brain stem tissue was enzymatically digested then mechanically dissociated with the gentleMACS Dissociater with Heaters. Debris Removal Solution and Red Blood Cell Removal Solution were then used to remove myelin, cell debris, and erythrocytes. Astrocytes were isolated using Anti-ACSA-2 MicroBeads. Microglia were removed by shaking in an incubator overnight and astrocytes were verified as negative for microglia through Iba1 staining. Isolated astrocytes were cultivated in DMEM/F12 with 10% FBS penicillin/streptomycin for two weeks before use.

#### Astrocyte Lipid droplets staining and quantification

Lipid droplets in astrocytes were stained and quantified as described^43^. HCS LipidTOX^TM^ Red Neutral Lipid Stain was used to stain astrocytes following the manufacturer’s instruction. Isolated astrocytes were cultivated in DMEM/F12 with 10% FBS penicillin/streptomycin for two weeks before use. Medium was removed and cells were washed with PBS 3 times. Cells were then fixed in 4% Paraformaldehyde for 30 minutes and stained with 1X LipidTOX^TM^ for 30 minutes at room temperature. A Zeiss LSM 880 with Airyscan was used to collect Z section images and generate three-dimensional images. The surface module of Imaris 9.3.0 was used to quantify the number and volume of lipid droplets from at least 3 animals or batches of cell cultures; n = 3–8 coverslips/condition; 8 ± 2 cells/coverslip. Each data point represents the averaged value of all cells on a coverslip.

#### Lenti-Ndst1shRNA transfection

Astrocytes isolated from E18 pregnant C57BL/6J, *APOE3* and *APOE4* knock-in mice were seeded on Poly-D-Lysine coated 12-well or 6-well plate with coverslips at a density of 3x10^5^ or 5x10^3^ well with complete medium (DMEM/F12 supplemented with 10% Fetal Bovine Serum and 1% Penicillin & Streptomycin), respectively. The lentivirus stocks (Lenti-Ndst1shRNA and scramble) were thawed on ice before transfection. The appropriate volume of virus was diluted into the media (DMEM/F12 supplemented with 10% FBS only) in order to achieve the desired MOI (Multiplicity of infection, 5-10 cells) of virus according to the stock concentration. After 48h incubation, the medium was aspirated, and the cells were washed once with PBS and incubated with complete medium for another 3 days. Astrocytes from 12-well plates were collected for RT-PCR and 6-well plates with coverslip were used for immunostaining. Results were obtained from 4 to 6 independent experiments performed in duplicate wells. Transduction efficiency was assessed by analyzing the number of cells that showed positive for GFP fluorescence after fixation and by staining all cell nuclei with DAPI.

#### RNA extraction and RT-qPCR from *Drosophila* and human cultured cells

Total RNA was isolated from cultured cells described above using TRIzol and PureLink RNA Mini Kit (Cat# 12183018A, Thermo Fisher). Extracted RNAs were reverse transcribed to cDNA using the SuperScript IV VILO Master (Cat# 11756050, Thermo Fisher). Relative mRNA levels of target genes (Ndst1, Plin2, Gfap, Fasn, Acc, Pparg, Cpt1α, Pgc1α) were amplified by SYBR Green real-time PCR assays in QuantStudio 6 System. The primer sequences are listed in Table 1. The relative gene expression level was calculated by the comparative Ct (ΔΔCt) method when compared to the internal control genes (Hprt, Gapdh, β-actin, RPL13) for individual samples. Results were obtained from 4 independent experiments performed in duplicate wells.

**Table 1.**
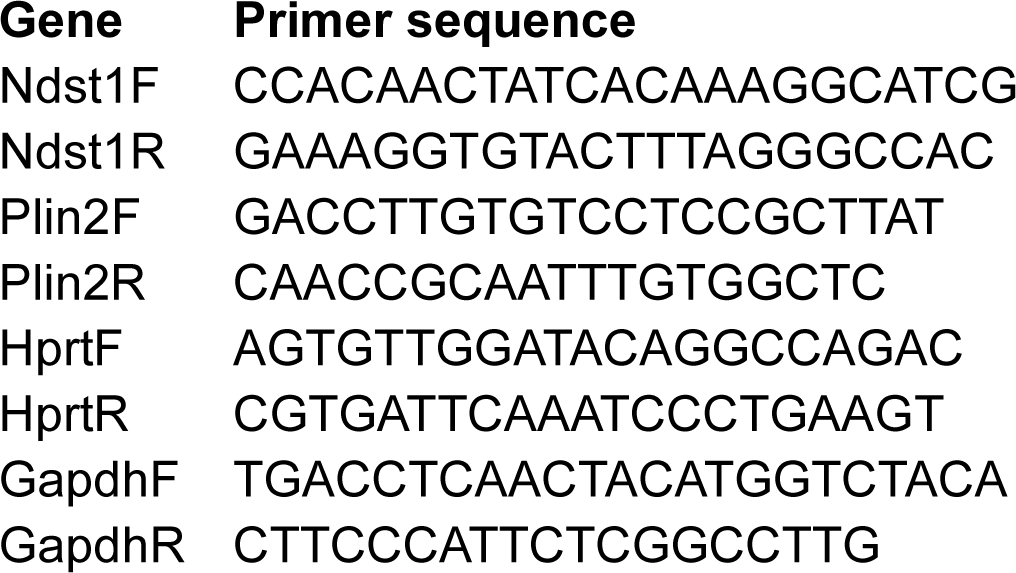
Primers sequences.

#### LipidTOX™ staining

Lipid droplet staining was performed according to the product manual (HCS LipidTOX™ Neutral Lipid Stains, Cat# H34476 HCS LipidTOX™ Red neutral lipid stain). Briefly, the incubation medium was aspirated, and enough formaldehyde fixative solution was added to a 6-well plate to cover cells for 30 minutes at room temperature. Then the fixed cells were rinsed with PBS 3 times and incubated with LipidTOX™ neutral lipid stain (1:200 dilution) at room temperature for 30 minutes. The cells were covered with mounting medium containing DAPI (Cat#H-2000, Vector Laboratories, Inc). Images were then captured with Zeiss Laser-Scanning Microscopy (LSM) 880 with Airyscan detector. Z stack images were captured to generate three-dimensional volumetric data. The number and volume of intracellular lipid droplets were quantified using the surface module of Imaris 9.8.0.

### QUANTIFICATION AND STATISTICAL ANALYSIS

#### Avizo image processing

Three-dimensional reconstructions of mitochondria were generated using Avizo software. Outlines of mitochondria were traced through all slices where the structure was visible and interpolated between slices using the built-in Avizo function set to the appropriate slice depth of 100 μm. Images were taken of each mitochondrial reconstruction at an approximately equal zoom for analysis. Quantitative data was obtained using the built-in measurement function of Avizo and analyzed.

Flow Cytometry statistics were analyzed using GraphPad Prism v10.0 (https://www.graphpad.com/) and other statistics were analyzed using Minitab (https://www.minitab.com/en-us/). Details of statistical tests, sample sizes, n, what n represents, and levels of significance can be found in figures and their corresponding figure legends.

### SUPPLEMENTAL ITEM TITLES AND LEGENDS

Differential Gene Expression of PsnRNAi Drosophila vs Control

Differential Gene Expression of SflRNAi Drosophila vs Control

Differential Gene Expression of PsnRNAi;SflRNAi Drosophila vs Control

Differential Gene Expression of EXT1 KO Hep3B Cells vs Control

---

**Supplemental Figure 1.**
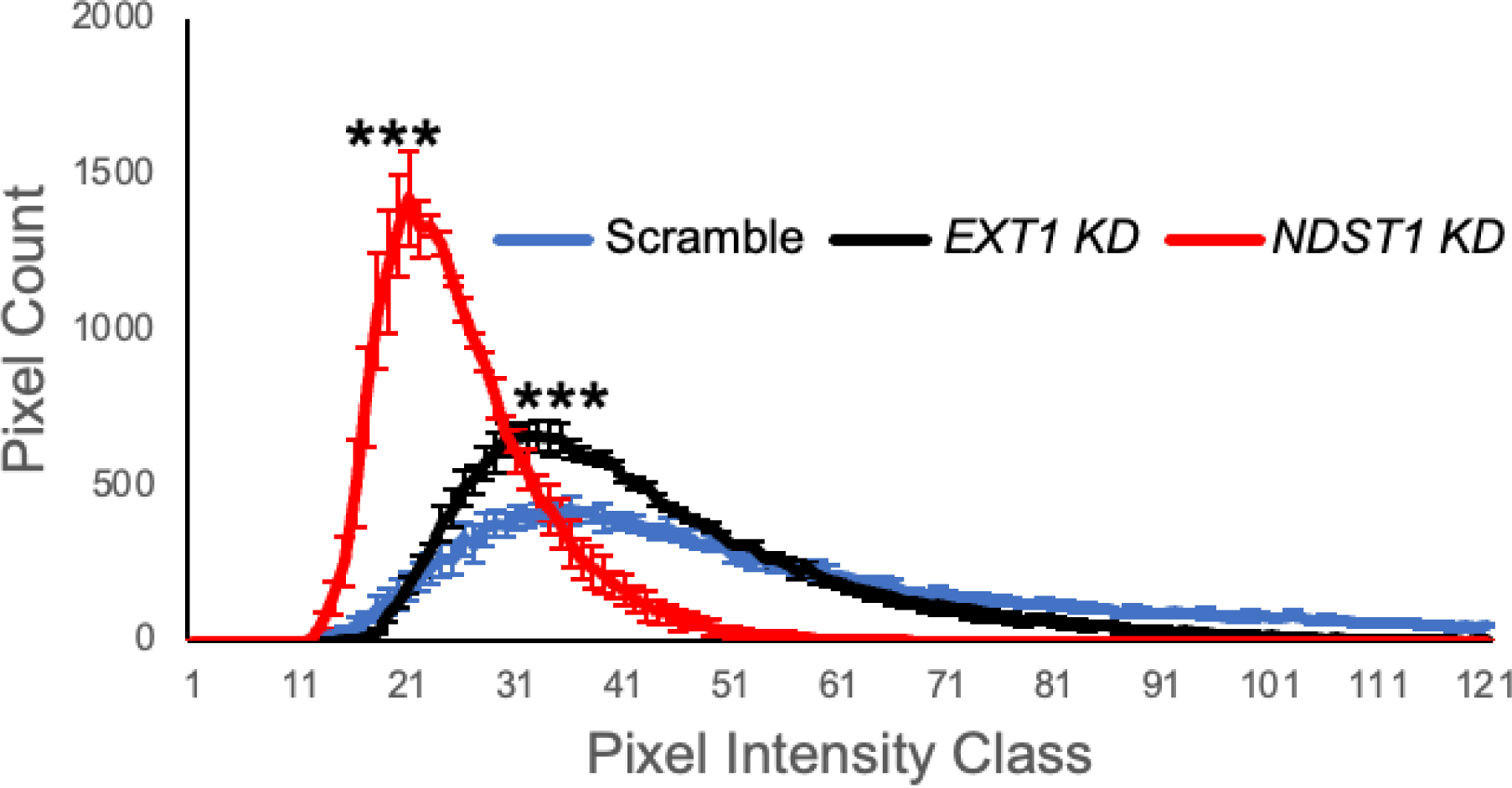
Histogram of LipidTOX^TM^ Red intensity distribution in HEK293 cells with knockdown of either *EXT1* or *NDST1*. Knockdown of NDST1 and EXT1 significantly decreases the pixel intensity of LipidTox Red detection of neutral lipid in HEK293T Cells (Kolmogorov-Smirnov Test, ***p<0.001).

**Supplemental Figure 2.**
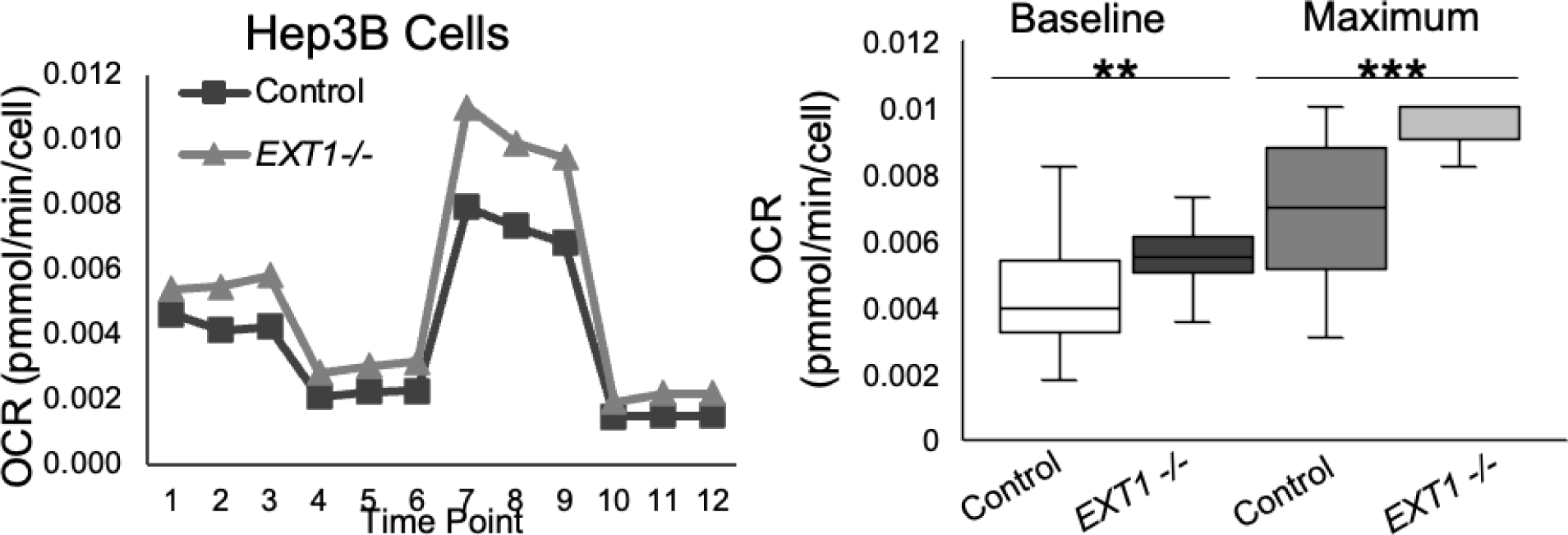
Oxygen consumption rate of *EXT1 -/-* Hep3B cells compared to wild type controls. Representative graph and quantification of Hep3B cells undergoing a Mito Stress Test using Seahorse Analyzer to quantitate oxygen consumption. Knockout of EXT1 significantly increases the Baseline and Maximal OCR of Hep3B cells [t test, n=24 (8 wells x 3 time points) per genotype, *p<0.05, **p<0.01, ***p<0.001]. Experiment was conducted in triplicate.

**Supplemental Table 1.** HEP3B *EXT1-/-* vs. Wild Type: Differential Gene Expression.

| Category/Gene | Log<sub>2</sub> Fold Change | P-adjusted value |
| --- | --- | --- |
| Cholesterol Biosynthesis |  |  |
| *HMGCR* | -0.95 | 5.2E-30 |
| *FDFT1* | -0.72 | 7.3E-42 |
| *SQLE* | -1.29 | 2.2E-87 |
| *SOAT1* | -0.66 | 4.4E-21 |
| *SREBF2* | -0.83 | 1.5E-41 |
| Lipid Transport |  |  |
| *APOE* | -1.04 | 4.5E-32 |
| *APOC1* | -1.16 | 1.4E-60 |
| *APOC2* | -1.45 | 2.6E-48 |
| *APOB* | -1.12 | 5.2E-59 |
| *MTTP* | -4.60 | 2.1E-153 |
| *APOA4* | 5.03 | 1.0E-14 |
| *ABCA7* | 1.22 | 3.0E-24 |
| *ABCA10* | 1.94 | 4.2E-5 |
| *ABCA12* | 2.88 | 0.0022 |
| *PCSK9* | -2.90 | 7.7E-91 |
| *MFSD2A* | -2.29 | 8.2E-12 |
| Autophagy |  |  |
| *SQSTM1* | 0.98 | 1.6E-98 |
| *ULK1* | 1.11 | 1.7E-61 |
| *PINK1* | 1.10 | 2.6E-51 |
| *ATG9B* | 1.49 | 0.0035 |
| *SRC* | 1.07 | 1.6E-40 |
| *PRKAG2* | 0.78 | 3.9E-6 |
| Mitochondrial Biogenesis |  |  |
| *PPARGC1A* | 0.898 | 2.1E-15 |
| *CHD9* | 0.99 | 1.6E-7 |
| *MAPK11* | 1.31 | 4.6E-13 |

**Supplemental Table 2.**
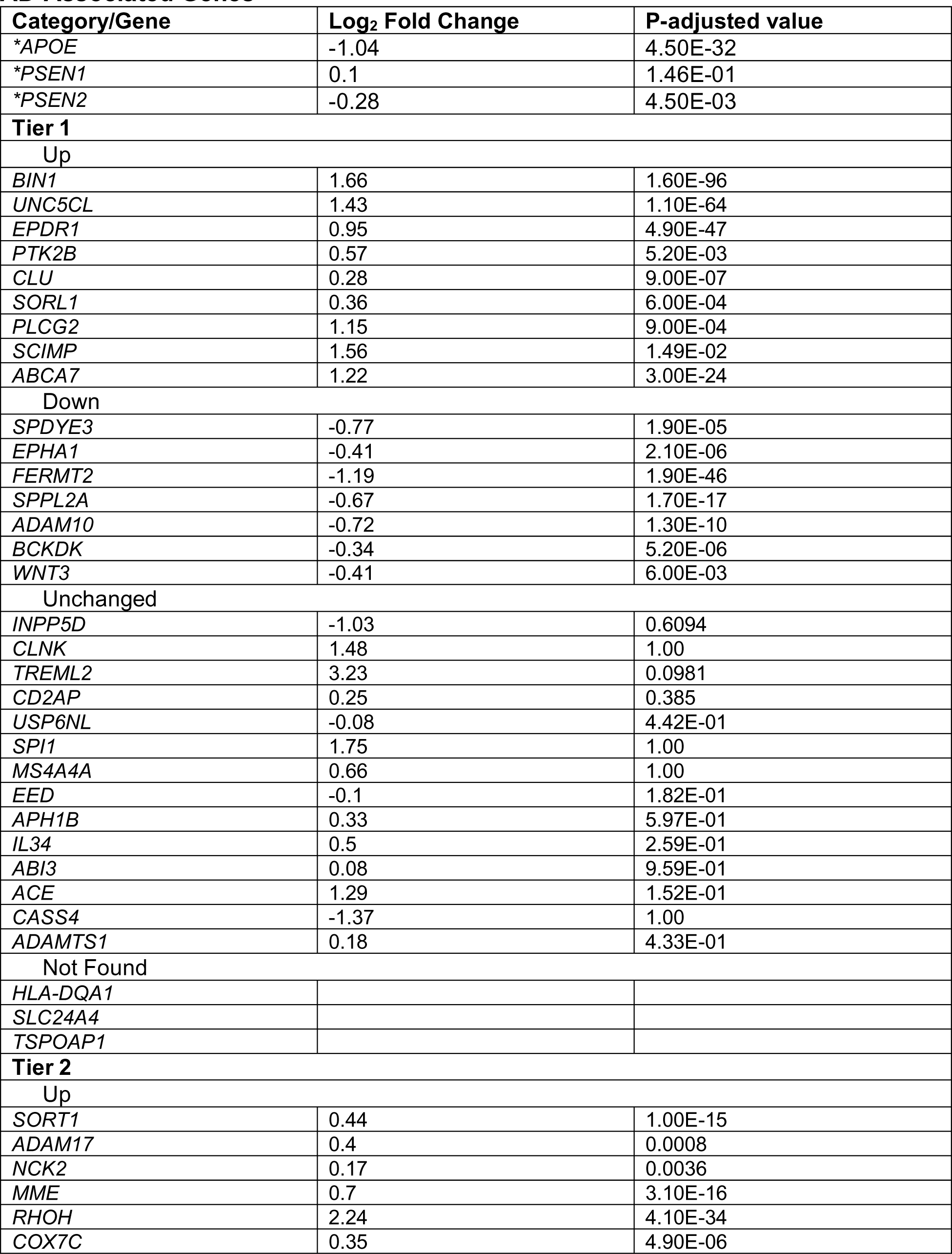
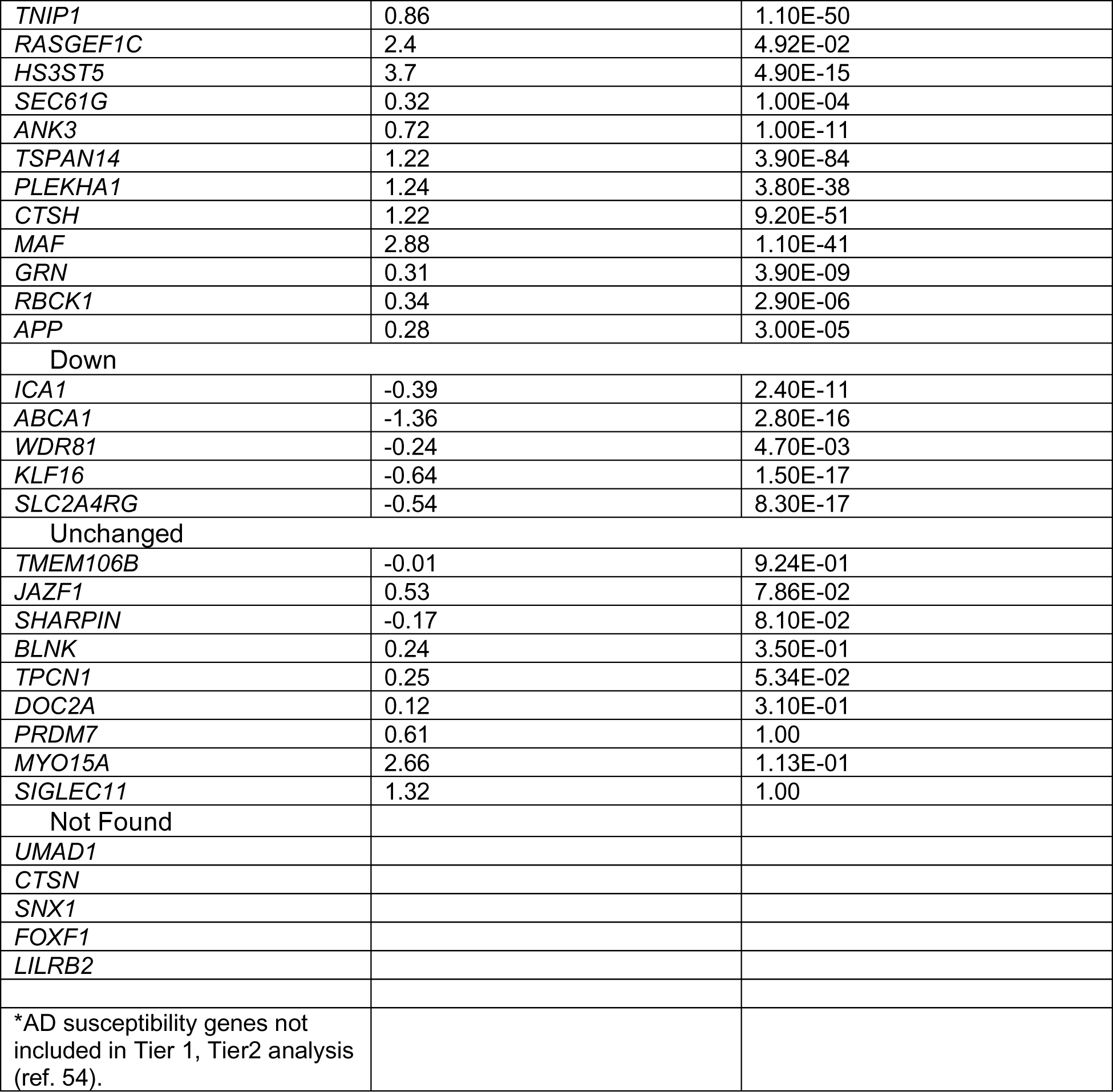
HEP3B *EXT1-/-* vs. Wild Type: Differential Gene Expression, AD-Associated Genes.

**Supplemental Table 3.** GSEA Sets with Upregulation in *EXT1-/-* HEP3B Cells (FDR< 0.25)

|  |  |
| --- | --- |
| **Autophagy** | **Mitochondria** |
| Selective Autophagy<br>Aggrephagy<br>Mitophagy | Activation of BAD and translocation to mitochondria<br>Intrinsic pathway for apoptosis |
| **Lipids** | **MAPK Signaling** |
| Biological Oxidations<br>Assembly of active LPL and LIPC lipase complexes | ERK/MAPK Targets<br>FLT3 Signaling<br>MAPK targets nuclear events mediated by MAP Kinases<br>FLT3 Signaling in disease<br>Signaling by MET<br>Nuclear events kinase and transcription factor activation<br>CREB1 phosphorylation through NMDA receptor-mediated activation of RAS signaling<br>Activation of SMO<br>FC epsilon receptor (FCERI) signaling<br>Activation of NMDA receptors and postsynaptic events |
| **Cytoskeleton** | **Inflammation** |
| Intraflagellar transport<br>Recycling pathway of L1<br>Formation of tubulin folding intermediates by CCT/TriC<br>L1CAM Interactions<br>Cilium Assembly<br>Crosslinking of collagen fibrils<br>CDC42 GTPase cycle<br>Elastic Fibre Formation<br>Molecules associated with elastic fibres<br>RAC1 GTPase cycle<br>Post-chaperonin tubulin folding pathway<br>Carboxyterminal post-translational modifications of tubulin<br>Translocation of SLC2A4 (GLUT4) to the plasma membrane | Interleukin 17 signaling<br>MHC Class II Antigen Presentation<br>Costimulation by the CD28 Family<br>Senescence associated secretory phenotype SASP<br>Interleukin 6 family signaling<br>Generation of second messenger molecules<br>FCERI mediated Ca<sup>2+</sup> mobilization<br>RHO GTPases activate NADPH oxidases<br>Signaling by interleukins<br>Pyroptosis |
| **ECM** | **Exocytosis** |
| GPVI mediated activation cascade<br>Extracellular matrix organization<br>Integrin cell surface interactions<br>ECM Proteoglycans<br>Netrin-1 Signaling<br>Degradation of the Extracellular Matrix<br>Assembly of collagen fibrils and other multimeric structures | Phase II conjugation of compounds<br>Regulation of gene expression in beta cells |
| **SARS-CoV-2** | **Miscellaneous** |
| Translation of SARS-CoV-2 Structural Proteins<br>SARS-CoV-2 Infection | Sulfonation of small molecules<br>Protein folding<br>Gap junction trafficking and regulation<br>G-alpha 12/13 signaling events<br>Organelle Biogenesis and Maintenance |

**Supplemental Table 4.** GSEA Sets with Downregulation in *EXT1-/-* HEP3B Cells (FDR < 0.25)

|  |  |
| --- | --- |
| **Autophagy** | **Aberrant MAPK Signaling** |
| Ubiquitin-specific processing proteases<br>Late endosomal microautophagy | Signaling by PDGFR in Disease<br>Signaling by BRAF and RAF1 Fusions |
| **Metabolism and Lipids** | **DNA Repair** |
| Diseases associated with glycosylation precursor biosynthesis<br>Glycogen storage diseases<br>Nicotinate metabolism<br>Metabolism of steroid hormones<br>Metabolism of fat-soluble vitamins<br>Plasma lipoprotein clearance | Formation of TC-NER Pre-Incision Complex<br>Formation of Incision Complex in GG-NER<br>Resolution of AP-sites via the multiple nucleotide patch replacement pathway<br>Dual incision in TC-NER<br>Base excision repair<br>Processing of DNA double strand break ends<br>Base excision repair AP site formation |
| **RNA Regulation** | **Cell Cycle Regulation and DNA Replication** |
| RNA Polymerase III Transcription Initiation from Type 1 Promoter<br>MicroRNA miRNA biogenesis<br>RNA Polymerase I Transcription<br>Regulation of mRNA stability by proteins that bind AU-rich elements<br>Attenuation Phase | Conversion from APC/C:Cdc20 to APC/C:CDdh1 in late anaphase<br>Transcription of E2F targets under negative control by DREAM Complex<br>Cyclin A CDK2 Associated Events at S-Phase Entry<br>Activation of the Pre-Replicative Complex<br>Polymerase switching on the C-strand of the telomere<br>DNA Replication |
| **HIV and Innate Immunity** | **Miscellaneous** |
| HIV Elongation arrest and recovery<br>HIV Transcription Initiation<br>Host Interactions of HIV-factors<br>HIV transcription elongation<br>Initial triggering of complement<br>Cytosolic sensors of pathogen-associated DNA | Disassembly of the destruction complex and recruitment of AXIN to the membrane<br>Meiotic Recombination |

**Supplemental Table 5:**
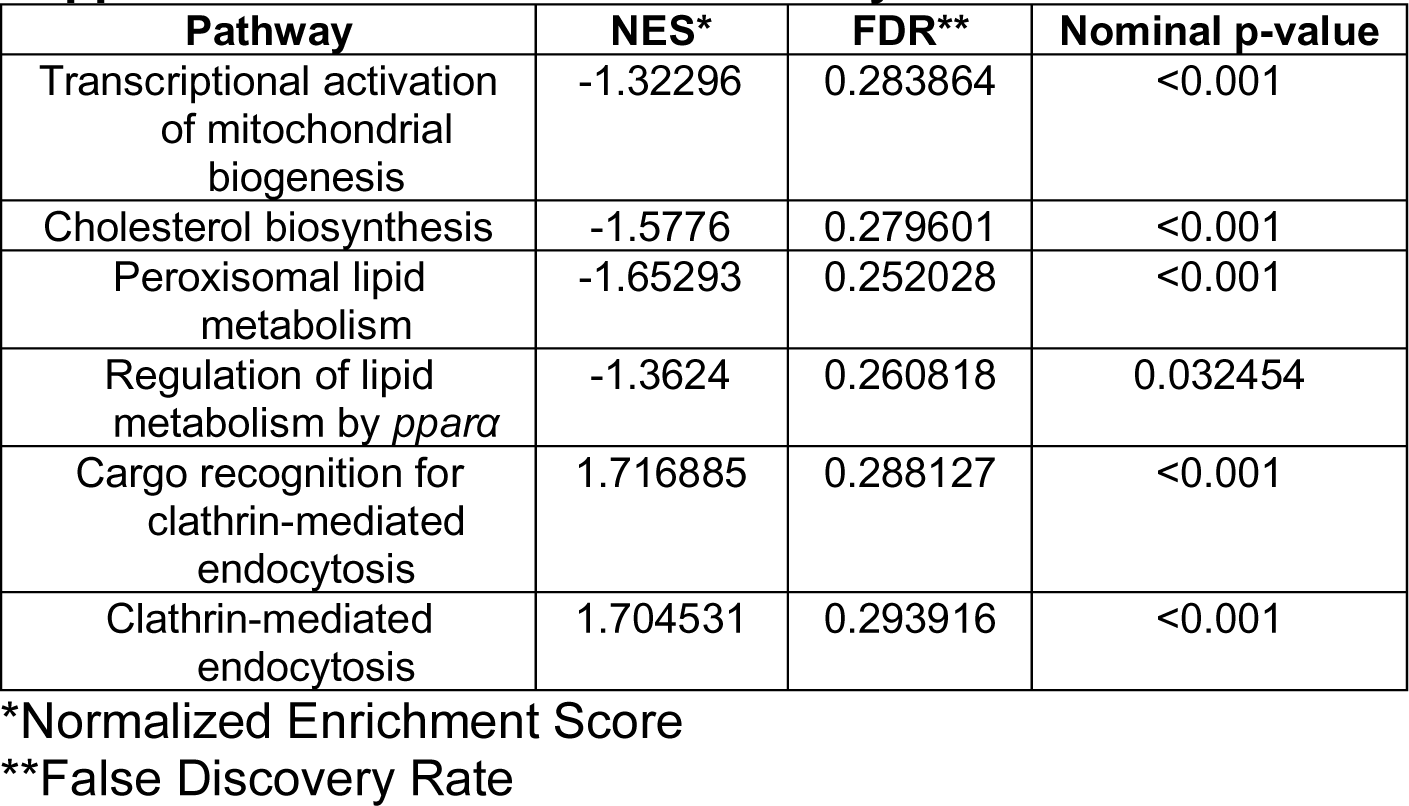
GSEA Pathways with 0.3 > FDR > 0.25.

